# High spatiotemporal resolution in vivo imaging reveals preferential PACAP38 action on dural vasculature and a functional therapeutic readout

**DOI:** 10.64898/2026.11.10.765588

**Authors:** Leonora Sperling, Caroline Degel, Kevin Zitelli, Jonathan Zapata, Douglas R. Ollerenshaw, Tanja Bruun, Taha Janjua, Franziska Wichern, Sarah Louise Christensen, Ayodeji A. Asuni, Jonathan Nassi, Benjamin J. Hall, Paolo Botta

## Abstract

PACAP38-induced cephalic vasodilation is used to assess functional PACAP pathway inhibition in humans, but the vascular populations underlying this response are not fully resolved. Defining these compartments in a preclinical model could help connect clinical pharmacodynamic responses with the underlying vascular mechanisms and support further development of PACAP-targeted therapies. Using chronic cranial windows and head-mounted microscopy, we tracked meningeal vascular responses in freely moving mice. Nitroglycerin, levcromakalim, and PACAP38 each produced net vasodilation, yet the underlying responses were markedly heterogeneous and distributed differently across the vascular populations. Notably, PACAP38 evoked greater early vasodilation in dural than pial vessels, and anti-PACAP pretreatment attenuated PACAP38-induced vasodilation. These findings show that net vasodilation can conceal vessel-level response patterns and identify preferential early dural vasodilation as a spatially defined, pharmacologically modifiable feature of the systemic PACAP38 response. This phenotype provides a translational platform for mechanistic interrogation and evaluation of functional pathway inhibition by migraine-targeted therapies.

## Introduction

Migraine is a highly prevalent neurological disorder, affecting an estimated 15.2% of the global population (Steinmetz et al., 2024). Migraine ranks among the leading causes of disability worldwide, yet the disorder remains underrecognized and frequently undertreated, and its underlying mechanisms are incompletely understood (Raggi et al., 2024; Steinmetz et al., 2024). Migraine is clinically heterogeneous, with substantial variation in attack frequency, aura status, associated symptoms and comorbidities, and response to therapy. This phenotypic diversity likely reflects heterogeneity in the underlying molecular and physiological mechanisms of migraine and complicates both mechanistic investigation and therapeutic development (Raggi et al., 2024). As such, extensive research has aimed to dissect migraine-relevant pathways, with pharmacological provocation studies serving as an important tool in both preclinical and clinical research (Guo et al., 2024; Olesen, 2024). Several established migraine-provoking agents engage distinct molecular pathways, including the nitric oxide donor nitroglycerin, the ATP-sensitive potassium (KATP) channel opener levcromakalim, and the neuropeptides calcitonin gene-related peptide (CGRP) and PACAP38 (Karsan et al., 2024; Olesen, 2024; Raffaelli et al., 2024). These agents can provoke migraine-like attacks in people with migraine that closely resemble spontaneous attacks, although induction rates vary across agents, study populations, and individuals (Al-Karagholi et al., 2019; Amin et al., 2014; Ashina et al., 2022; Onderwater et al., 2020). Their ability to produce attacks resembling spontaneous migraine supports the use of provocation models to interrogate migraine-relevant signaling pathways.

Within provocation paradigms, measurable physiological responses are particularly valuable given the lack of an established objective biomarker for migraine (Olesen, 2024). The close anatomical and functional coupling between trigeminal afferents and the meningeal vasculature makes vascular dynamics a useful readout of migraine-relevant trigeminovascular signaling (Arquizan et al., 2025; Wiggers et al., 2022). Accordingly, changes in meningeal vessel caliber have been used experimentally to monitor the vascular responses to migraine-relevant pathway activation, although dilation of the larger arteries assessed by human Magnetic Resonance Angiography (MRA) does not consistently accompany migraine attacks and cannot, by itself, establish that an attack has occurred (Arquizan et al., 2025; Schoonman et al., 2008; Wiggers et al., 2022). Though intracranial pain sensitivity has traditionally been attributed predominantly to the dura, mechanical stimulation of the pia mater and small cerebral vessels has been shown to evoke pain in awake patients, supporting examination of vascular responses across both dural and pial compartments (Fontaine et al., 2018).

PACAP is an endogenous neuropeptide directly implicated in human migraine and an actively pursued therapeutic target (Ashina et al., 2024; Pellesi et al., 2024). PACAP signaling is implicated in migraine-relevant trigeminovascular biology, in which trigeminal sensory afferents closely appose meningeal blood vessels – an anatomical arrangement that supports bidirectional coupling between vascular dynamics and nociceptive signaling (Ashina et al., 2019; Pellesi et al., 2024). PACAP38, the 38-amino-acid form of PACAP, is a potent vasodilator that produces sustained arterial dilation in human provocation studies (Amin et al., 2012; Arquizan et al., 2025). Notably, superficial temporal artery (STA) dilation served as the primary pharmacodynamic endpoint in a human proof-of-mechanism study (Rasmussen et al., 2023). This illustrates the translational utility of provoked vascular responses as functional readouts of PACAP pathway inhibition, yet the spatial organization of PACAP38-induced vascular responses within meningeal circulation remains incompletely resolved.

Despite the value of meningeal vascular dynamics as a physiological readout, methodological limitations continue to constrain cross-species comparisons and the translation of preclinical findings into clinical applications that benefit patients. Much preclinical work has relied on anesthetized rodent preparations, despite evidence that anesthesia substantially alters cerebrovascular tone and blood flow (Rakymzhan et al., 2021; Tran & Gordon, 2015; Wen et al., 2026). Meningeal studies have also frequently used acute or terminal preparations with surgically exposed meninges which can yield different sensory responses than those observed in awake animals (Blaeser et al., 2024). Awake head-fixed imaging avoids anesthesia but constrains the animal, which may limit observation of vascular responses during natural behaviors and pharmacological treatment. Although habituation can reduce stress associated with restraint (Witztum et al., 2023), freely moving preparations provide an opportunity to measure meningeal vascular dynamics in a less constrained behavioral context. Other mechanistic studies have examined isolated vessels ex vivo, providing precise pharmacological information but removing the vessels from their intact neural and physiological environment (Syed et al., 2012). In parallel, vascular approaches based on regional blood flow signals or measurements of major cranial arteries provide limited information about the dynamics of defined vessel segments within the broader meningeal network (Arquizan et al., 2025; Sokolov et al., 2021). Recent miniature fluorescence microscopy approaches have enabled stable meningeal vessel imaging in freely moving mice (Degel et al., 2024), providing an opportunity to examine pharmacologically induced vascular dynamics under more physiological conditions.

Here, we combined chronic cranial windows with head-mounted miniature one-photon microscopy to track individual meningeal vessel segments in non-anesthetized, freely moving mice. We used vascular diameter dynamics as a quantifiable physiological readout of migraine-relevant pathway engagement rather than as a surrogate for migraine itself. We first tested if the platform could resolve responses to the mechanistically distinct migraine provocateurs nitroglycerin, levcromakalim, and PACAP38. We then examined if vessel-level analysis revealed biologically organized heterogeneity, with particular emphasis on the spatial organization of the PACAP38 response. Finally, we tested if an anti-PACAP monoclonal antibody attenuated the PACAP-dependent vascular phenotype, thereby evaluating the model as a functional in vivo readout of pathway inhibition and as a platform for mechanistic and translational therapeutic investigation.

## Results

### Vasoactive migraine provocation agents increase meningeal vessel diameter in freely moving mice

To determine whether meningeal vascular responses could be resolved longitudinally in freely moving mice, we combined chronic cranial windows with miniature fluorescence microscopy as previously described (Degel et al., 2024). This approach enabled quantitative, vessel-resolved imaging of meningeal vascular dynamics during pharmacological challenge, allowing diameter changes in defined vessel segments to be tracked throughout each imaging session.

We first optimized the optical preparation for reliable visualization of the meningeal vasculature. Thinned-skull preparations provided less reliable visualization of smaller vessels under our imaging conditions, particularly when the imaging region overlapped with cranial sutures. We therefore used chronic cranial-window preparation for subsequent experiments. Following cranial-window surgery and miniscope baseplate implantation, mice were allowed to recover for at least 4 weeks before imaging. On each experimental day, FITC-dextran was administered intravenously before a 15-min baseline recording, followed by pharmacological challenge and up to three 10-min post-injection imaging periods separated by 5-min pauses (Fig. 1a and more detailed in Fig. S1a).

**Figure 1.**
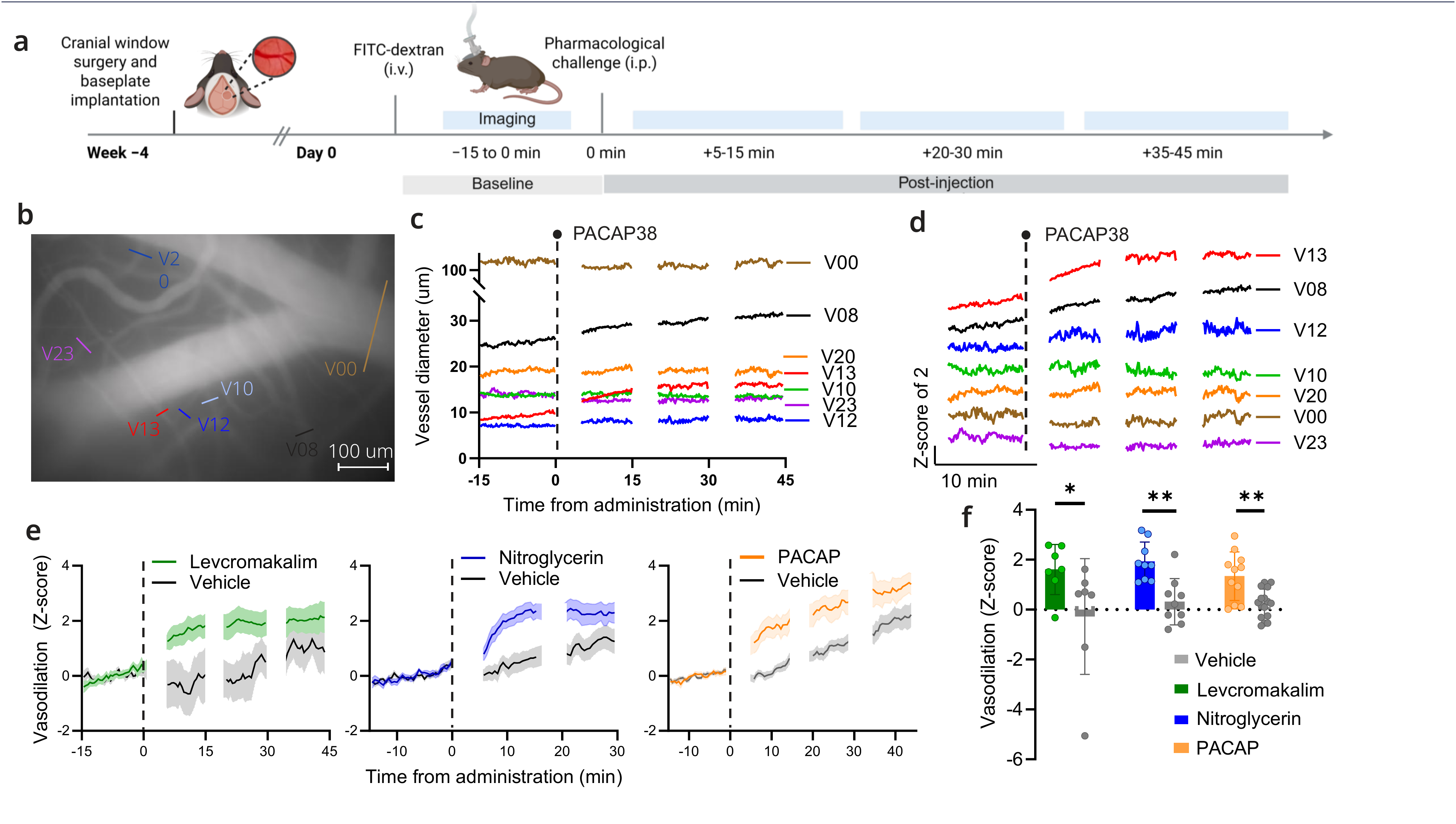
Levcromakalim, nitroglycerin and PACAP38 induce meningeal vasodilation. a, Experimental timeline. Mice underwent cranial-window surgery and baseplate implantation 4 weeks before imaging. On the imaging day, intravenous FITC–dextran administration was followed by a 15-min baseline, intraperitoneal injection of vehicle or vasoactive compound, and three 10-min imaging periods separated by 5-min intervals. Levcromakalim, nitroglycerin or PACAP38 were administered at 1, 10 and 0.1 mg/kg, respectively. Created in https://BioRender.com. **b,** Representative meningeal vascular field from a PACAP38-treated mouse showing seven tracked vessel segments. The vessel numbers were assigned sequentially according to the order in which the vessels were annotated. In this representative mouse, 67 vessels were annotated; therefore, V23 refers to the 23rd vessel annotated. Colored lines indicate locations at which vessel diameter was estimated. Mean image of entire imaging session. Scale bar, 100 µm. **c,** Diameter traces of the vessel segments shown in **b** before and after PACAP38 administration. The vertical dashed line indicates injection. **d,** Responses of the same vessels expressed as z-scores relative to the 15-min baseline. Traces are vertically offset by two z-score units for visualization. **e,** Time courses of vascular responses following administration of each compound and its study-matched vehicle. Responses are expressed as z-scores relative to the 15-min baseline. Vessel-level responses were averaged within mouse at each time point and then across mice within treatment groups. Lines show the mean and shaded regions indicate s.e.m. The dotted line indicates compound administration occurred at 0 min. Gaps in traces correspond to 5-min pauses between imaging periods. Vehicle is in gray while levcromakalim in green, nitroglycerin in blue and PACAP in orange. **f,** Mean vasodilation during the first 10 min after administration of each compound and its study-matched vehicle (same color code as in panel e). Vessel-level responses were averaged over the predefined response period and within each animal/session to generate one response value per animal/session. Points represent animal/session means. Levcromakalim: n = 7 mice (total of 179 vessels); vehicle, n = 7 mice (total of 134 vessels). Nitroglycerin: n = 9 mice (total of 317 vessels); vehicle, n = 9 mice (total of 336 vessels). PACAP38: n = 11 mice (total of 324 vessels); vehicle, n = 16 mice (total of 496 vessels). Levcromakalim (1 mg/kg) and PACAP38 (0.1 mg/kg) were compared with their respective vehicles using two-sided Mann–Whitney U tests (levcromakalim: U = 41, P = 0.0379; PACAP38: U = 141, P = 0.00958). Nitroglycerin (10 mg/kg) and vehicle were compared within the same mice using a two-sided Wilcoxon matched-pairs signed-rank test (W = 0, P = 0.00391). Each comparison represented a separate prespecified study-level hypothesis; no correction was applied across studies. *P < 0.05; **P < 0.01. Bars show mean ± s.e.m.

The cranial window was positioned to provide optical access to the middle meningeal artery and surrounding meningeal vasculature. Individual meningeal vessels remained stably visible throughout the approximately 1h imaging session, permitting vessel-specific vascular dynamics to be tracked over time in freely moving animals. On average, 53.4 ± 8.2 vessels per mouse remained identifiable throughout the session (mean ± SEM; n = 48 mice). Figure 1b shows a mean image generated across an entire PACAP38 challenge session, with individual vessel segments selected for diameter analysis, while Figure 1c shows the corresponding diameter traces over time. Vessel diameter was quantified independently for each annotated vessel segment, preserving vessel-level information for subsequent analysis of pharmacologically induced vascular responses. Vessel diameters varied substantially across the imaged field (Fig. 1b-c), and responses were therefore normalized to each vessel’s own baseline mean and variability to generate vessel-specific z-scores (Fig. 1d). The normalized traces revealed marked vessel-to-vessel heterogeneity together with an overall tendency toward vasodilation in response to PACAP38 over time.

To determine whether the imaging approach detected pharmacologically induced changes in meningeal vessel diameter, mice were given the migraine-provoking agents levcromakalim (1 mg/kg), nitroglycerin (10 mg/kg), or PACAP38 (0.1 mg/kg), intraperitoneally (see Methods for dosing rationale).

Time-resolved responses showed a clear vasodilatory shift for each migraine-provoking agent relative to its study-matched vehicle (Fig. 1e). To quantify these responses across migraine-provoking agents and their respective vehicles, we evaluated mean vasodilation during the first 10 min following administration (Fig. 1f). Across studies, all three migraine-provoking agents produced greater mean vasodilatory responses than their corresponding vehicle treatments. Levcromakalim increased meningeal vasodilation relative to its study-matched vehicle (Mann– Whitney U = 41, P = 0.0379). Nitroglycerin similarly increased vasodilation compared with vehicle in the same mice tested (Wilcoxon matched-pairs signed-rank test, W = 0, P = 0.00391). PACAP38 also increased meningeal vasodilation relative to its study-matched vehicle (Mann–Whitney U = 141, P = 0.00958). The magnitude of the normalized response was broadly similar across levcromakalim (1.61 ± 0.38 z-score units, n = 7 mice), nitroglycerin (1.93 ± 0.26 z-score units, n = 9 mice), and PACAP38 (1.34 ± 0.29 z-score units, n = 11 mice). Group means expressed as z-scores and percent change from baseline are summarized in Supplementary Table 1. Together, these findings show that established migraine-provoking agents increase meningeal vessel diameter and that these responses can be quantified in freely moving mice.

### PACAP38 preferentially dilates small meningeal vessels, whereas size-dependent responses to the other agents are less consistent

Although the three migraine-provoking agents produced broadly similar mean vasodilatory responses when responses were averaged across vessels, individual vessel responses varied markedly in both magnitude and direction (Fig. 2a): 50.3% of levcromakalim-treated vessels, 53.3% of nitroglycerin-treated vessels, and 45.0% of PACAP38-treated vessels were classified as vasodilating during the first 10 min after intraperitoneal administration, whereas 6.7%, 4.7% and 5.9% of vessels, respectively, were classified as vasoconstricting. Thus, the net vasodilatory response reflected heterogeneous responses across individual vessels rather than uniform dilation throughout the vascular network.

**Figure 2.**
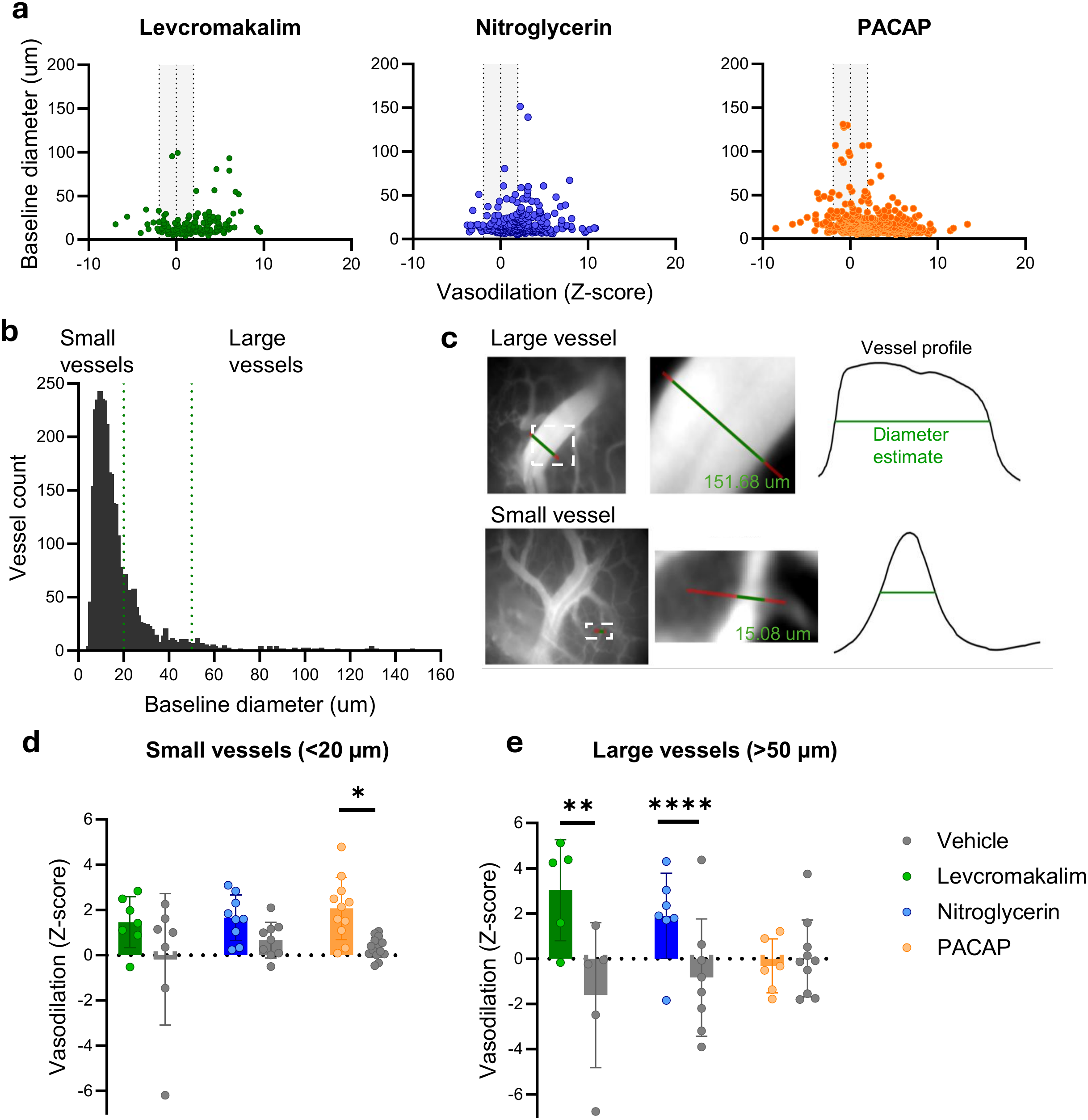
Vessel size contributes to heterogeneity in meningeal vascular responses to migraine-provoking agents. **a,** Relationship between vasodilation during the first 10 min after administration and baseline vessel diameter for levcromakalim (1 mg/kg, green), nitroglycerin (10 mg/kg, blue), and PACAP38 (0.1 mg/kg, orange). Each point represents one accepted vessel. The central vertical dashed line indicates z = 0, corresponding to no change from baseline, while the two flanking dashed lines at z = ±1.96 indicate thresholds for responses exceeding 1.96 standard deviations from the vessel-specific baseline mean; negative and positive values indicate vasoconstriction and vasodilation, respectively. **b,** Distribution of baseline diameters across all accepted vessels. Dashed lines indicate the thresholds used to define small vessels (<20 µm) and large vessels (>50 µm); vessels with baseline diameters of 20–50 µm were excluded from the size-stratified analyses. **c,** Representative large and small vessel segments and corresponding transverse pixel-intensity profiles used for diameter estimation. Green lines indicate the detected vessel diameters (151.68 and 15.08 µm, respectively). **d,** Mean vasodilation during the first 10 min after compound or study-matched vehicle administration in small vessels (<20 µm). Vessel-level z-scores were averaged within each animal/session. Bars show mean ± s.e.m.; points represent animal/session means. Levcromakalim: n = 7 mice (total of 136 vessels); vehicle, n = 7 mice (total of 93 vessels). Nitroglycerin: n = 9 mice (total of 241 vessels); vehicle, n = 9 mice (total of 252 vessels). PACAP38: n = 11 mice (total of 258 vessels); vehicle, n = 15 mice (total of 369 vessels). Planned treatment-versus-vehicle contrasts: levcromakalim, P = 0.144; nitroglycerin, P = 0.0948; PACAP38, P = 0.0161. **e,** Mean vasodilation during the first 10 min after compound or study-matched vehicle administration in large vessels (>50 µm). Vessel-level z-scores were averaged within each animal/session. Bars show mean ± s.e.m.; points represent animal/session means. Levcromakalim: n = 5 mice (total of 9 vessels); vehicle, n = 5 mice (total of 7 vessels). Nitroglycerin: n = 7 mice (total of 12 vessels); vehicle, n = 8 mice (total of 13 vessels). PACAP38: n = 6 mice (total of 13 vessels); vehicle, n = 10 mice (total of 23 vessels). Planned treatment-versus-vehicle contrasts: levcromakalim, P = 0.00389; nitroglycerin, P = 1.30 × 10⁻ ; PACAP38, P = 0.585. Study-specific linear mixed-effects models included treatment, vessel-size category and their interaction as fixed effects, with mouse identity as a random intercept. The treatment × vessel-size interaction was not significant for levcromakalim (Wald χ²(1) = 3.61, P = 0.0575) but was significant for nitroglycerin (Wald χ²(1) = 5.01, P = 0.0252) and PACAP38 (Wald χ²(1) = 4.34, P = 0.0372). Planned treatment-versus-vehicle contrasts within each size category were Holm-adjusted across the small- and large-vessel comparisons within each study. **ns**, P ≥ 0.05; *P < 0.05; **P < 0.01; ****P < 0.0001.

Interestingly, both dilatory and constrictive responses were observed across the range of baseline vessel diameters following levcromakalim (1 mg/kg), nitroglycerin (10 mg/kg), and PACAP38 (0.1 mg/kg). To test whether this overall response masked size-dependent differences, we examined responses according to baseline vessel diameter. Baseline vessel diameters spanned a broad range across the dataset, from 4.5 to 151.5 µm (Fig. 2b). We initially tested whether the vascular response was directly correlated with baseline vessel diameter. PACAP38 responses decreased with increasing vessel diameter, consistent with stronger vasodilation in smaller vessels (slope = −0.0237 z-score/µm, Holm-adjusted *P* = 0.005). By contrast, in nitroglycerin and levcromakalim provocation, larger vessels tended to exhibit greater vasodilation, although these associations did not remain significant after correction (Holm-adjusted *P* = 0.0893 and 0.1396, respectively). Next, vessels were classified as small (<20 µm) and large (>50 µm) vessels, focusing on opposite ends of the vessel-size distribution. Representative examples of small and large vessels, together with the corresponding intensity profiles used to estimate vessel diameter, are shown in Fig. 2c. For levcromakalim, the effect of treatment tended to differ between small and large vessels, although this did not reach the conventional threshold for statistical significance (treatment x vessel-size interaction: Wald χ²(1) = 3.61, P = 0.0575). However, planned contrasts showed that levcromakalim significantly increased responses relative to vehicle in large vessels (Holm-adjusted *P* = 0.00389), but not in small vessels (*P* = 0.144, Fig. 2d, e). For nitroglycerin, the treatment effect differed significantly between small and large vessels (treatment x vessel-size interaction: Wald χ²(1) = 5.01, P = 0.0252), with a significant effect in large vessels (Holm-adjusted *P* = 1.30 × 10⁻□ ) but not in small vessels (*P* = 0.0948; Fig. 2d, e). PACAP38 showed the opposite size-dependent response pattern. The treatment effect differed significantly between small and large vessels (treatment x size interaction: Wald χ²(1) = 4.34, P = 0.0372), with a significant treatment effect in small vessels (Holm-adjusted P = 0.0161) but not in large vessels (P = 0.585; Fig. 2d, e).

Because dural vessels were generally smaller than pial vessels, we next examined whether PACAP38 responses also differed between these anatomical territories. This direct comparison provides a more biologically informative assessment of the response pattern than vessel caliber alone.

### PACAP38 preferentially evokes vasodilation in dural meningeal vessels

Having characterized the vascular response profiles of all three migraine-provoking agents, we focused subsequent analyses on PACAP38 because of its direct relevance to migraine biology, its therapeutic relevance, and its distinct vascular response profile. Because dural and pial vascular compartments differ in vascular architecture and caliber, and the dura represents a key trigeminovascular nociceptive compartment, we asked whether the apparent size dependence of the PACAP38 response reflected compartment-specific vascular responsiveness.

To distinguish dural from pial vessels within the meningeal vascular preparation, we first examined the organization of the two vascular compartments using complementary histological and imaging approaches. CD31 (PECAM-1), an endothelial cell marker that delineates the vascular endothelium and thereby identifies blood vessels, and von Willebrand factor labeling visualized the meningeal vasculature in the coronal plane, illustrating the superficial position of the meningeal vascular network relative to the underlying brain parenchyma (Fig. 3a). CD31 immunostaining of isolated dura mater revealed a dense vascular network containing relatively long, linear vessel segments that frequently formed small loops (Fig. 3b). In parallel, ex vivo one-photon miniscope imaging of WGA-labeled cortical preparations visualized the pial vasculature at the brain surface (Fig. 3c). Compared with the dural network, pial vessels (Fig. 3c) showed a distinct branching organization, with some vessel segments terminating abruptly within the imaging plane, consistent with penetrating arteries diving in to the cortex (Agarwal & Carare, 2021). These anatomical differences were summarized schematically, using BioRender’s AI tool, to illustrate the relative organization of dural and pial vessels in the *xy* and *z* planes (Fig. 3d). Using these features, each annotated vessel across the complete imaging dataset was individually reviewed (n=2452 vessels) and manually classified as dural, pial, or unknown. Vessels were retained as unknown when compartment identity could not be assigned with confidence.

**Figure 3.**
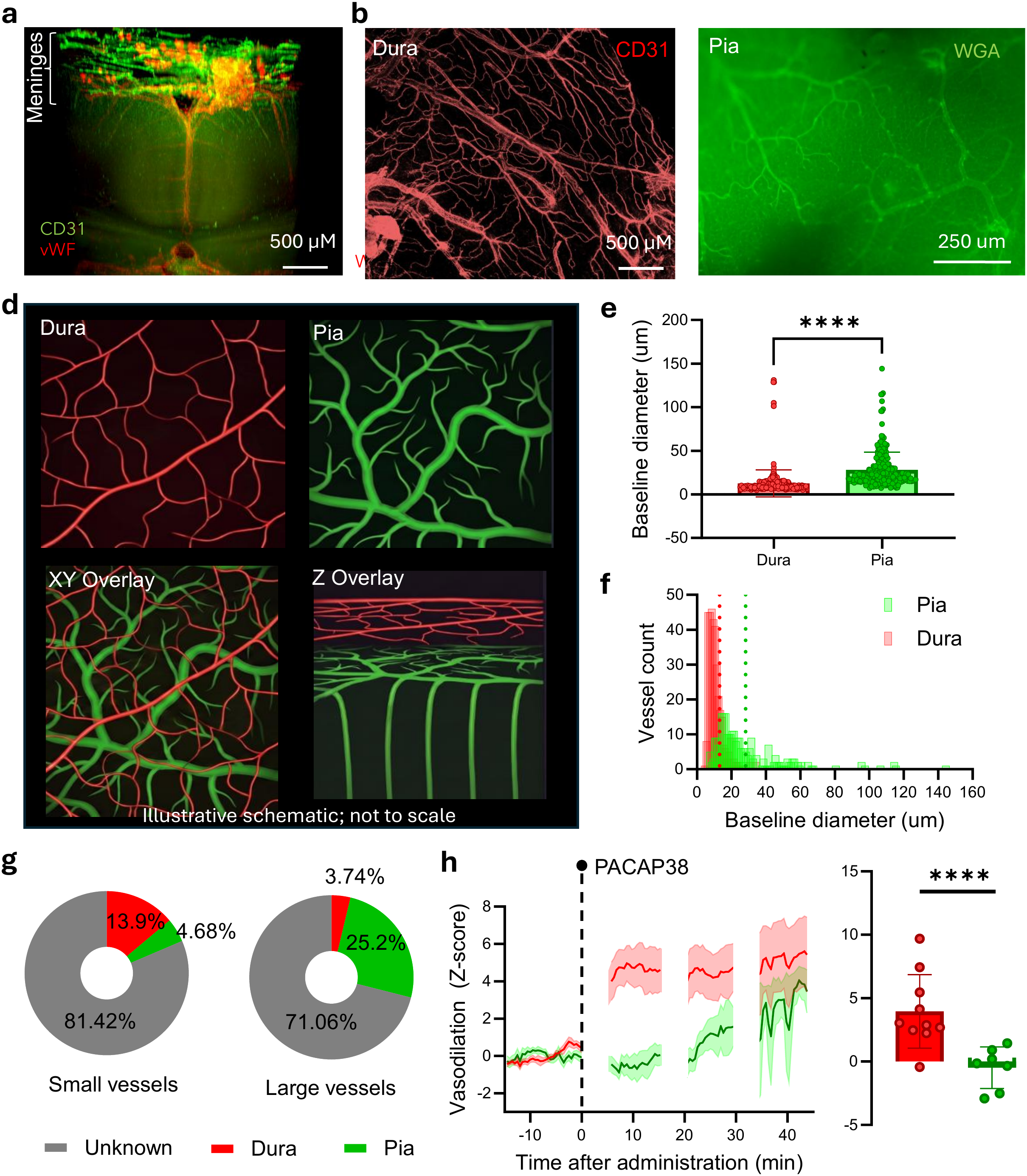
PACAP38 evokes stronger vasodilation in dural than pial vessels. **a,** Representative light-sheet fluorescence microscopy of an iDISCO-cleared preparation showing the meningeal vasculature labeled for CD31 and von Willebrand factor (vWF). Scale bar, 500 µm. **b,** Representative immunofluorescence image of isolated dura mater stained for the endothelial marker CD31. Scale bar, 500 µm. **c,** Representative one-photon miniscope image of pial vessels on the cortical surface following vascular labeling with Alexa Fluor 488-conjugated WGA. **d,** Illustrative schematic of the spatial organization of dural and pial vessels and their appearance in the *xy* and *z* planes. Dural vessels are shown in red and pial vessels in green. Generated with the assistance of generative AI using representative dural and pial histological images as visual references and subsequently assembled in BioRender. **e,** Baseline diameter of individual dural and pial vessels. Each point represents one accepted vessel. Dural vessels had a mean baseline diameter of 12.36 µm and pial vessels 28.54 µm. The analysis included 274 dural and 209 pial vessels from 25 mice across 44 imaging sessions. Statistical analysis: linear mixed-effects model with vascular compartment and study/source as fixed effects and mouse and imaging session as random intercepts; adjusted pia − dura = 17.02 µm, *P* = 2.16 × 10^-30^. **f,** Distribution of baseline vessel diameters in dural and pial compartments. Red and green histograms indicate dural and pial vessels, respectively. Dashed vertical lines indicate the mean baseline diameter for dural (12.36 µm) and pial (28.54 µm) vessels. **g,** Classification of small and large vessels as dural, pial, or unknown across imaging data set. Among small vessels, 265/1902 (13.9%) were classified as dural, 89/1902 (4.7%) as pial, and 1548/1902 (81.4%) as unknown. Among large vessels, 4/107 (3.7%) were classified as dural, 27/107 (25.2%) as pial, and 76/107 (71.0%) as unknown. Unknown vessels could not be confidently assigned to either the dural or pial compartment. **h, Left,** Time courses of PACAP38-induced vascular responses in dural (red) and pial (green) vessels. Responses are expressed as z-scores relative to the 15-min baseline. The dashed line indicates PACAP38 administration at 0 min; gaps correspond to 5-min pauses between imaging periods. Lines show the mean and shaded regions indicate s.e.m. **Right,** Mean PACAP38-induced vasodilation during the first 10-min post-injection imaging period in dural and pial vessels. Dura, n = 10 mice; pia, n = 7 mice; 13 unique mice were included, with 4 mice contributing measurements from both compartments. A maximum-likelihood linear mixed-effects model included vascular compartment as a fixed effect and mouse identity as a random intercept. The model coefficient was defined as pia minus dura (estimate = −4.579 z-score units, *P* = 2.20 × 10⁻□ ). Bars show mean ± s.e.m.; points represent mouse-level mean responses. \*\*\*\**P* < 0.0001.

Baseline vessel diameter differed markedly between the two compartments, with morphologically classified dural vessels averaging 12.36 µm and pial vessels 28.54 µm in diameter (274 dural and 209 pial vessels; Fig. 3e, f). A vessel-level linear mixed-effects model accounting for clustering by mouse and imaging session and adjusting for study/source estimated pial vessels to be 17.02 µm larger than dural vessels (n=274 dural and 209 pial vessels from 25 mice across 44 imaging sessions; P = 2.16 × 10⁻³ ). Consistent with this difference, dural vessels were more frequently represented among small vessels, whereas pial vessels were relatively more common among large vessels (Fig. 3g). As classification was intentionally conservative, 81.4% of small vessels and 71.0% of large vessels remained unclassified. Of all small vessels, 13.9% were classified as dural and 4.7% as pial, whereas among large vessels, 3.7% were classified as dural and 25.2% as pial.

We then examined whether PACAP38-induced vascular responses differed between dural and pial vessels. Time-resolved responses showed greater vasodilation in dural than pial vessels after PACAP38 administration (Fig. 3h). During the first post-injection response window, PACAP38 (0.1 mg/kg) was followed by a dural-vessel response significantly above baseline (mean = 3.961 z-score units; two-sided one-sample *t*-test, *t*(9) = 4.33, Holm-adjusted *P* = 0.00382), whereas the mean pial-vessel response was not significantly different from baseline (mean = −0.481 z-score units; *t*(6) = −0.77, Holm-adjusted *P* = 0.4684). Consistent with this, in the PACAP38-versus-vehicle comparison, the treatment × compartment interaction was significant (Wald χ²(1) = 11.134; *P* = 0.000848). Planned contrasts estimated a PACAP38-versus-vehicle difference of 3.652 z-score units in dura (Holm-adjusted *P* = 0.000119) and −0.765 z-score units in pia (Holm-adjusted *P* = 0.4663), indicating preferential vasodilation of dural over pial vessels. No clear compartment-specific pattern emerged for nitroglycerin, whereas levcromakalim tended towards the pial vessels driving the vasodilation, but a significant difference was not detectable (Fig. S2). Because dural and pial vessels also differed in baseline caliber, we next asked whether vessel diameter itself explained the compartment-specific PACAP38 response. Within the dural compartment, PACAP38 response was not associated with baseline vessel diameter (53 vessels from 10 mice; slope = 0.0096 z-score units/µm, Holm-adjusted P = 1.000). Similarly, no relationship was detected within pial vessels (37 vessels from 7 mice; slope = 0.0024 z-score units/µm, Holm-adjusted P = 1.000). Thus, the inverse relationship between vessel diameter and PACAP38 responsiveness observed across the combined dataset was not reproduced within either compartment, suggesting that the preferential dural response to PACAP38 is not readily explained by the smaller caliber of dural vessels.

Together, the functional imaging data indicate that systemic PACAP38 preferentially evokes early vasodilation in the dural vascular compartment.

### Anti-PACAP monoclonal antibody pretreatment attenuates PACAP38-induced dural vasodilation

We then tested whether PACAP neutralization could attenuate PACAP38-induced meningeal vasodilation. Mice were pretreated with an anti-PACAP monoclonal antibody (mAb, 30 mg/kg) or vehicle 24 hours before PACAP38 administration (Fig S1b; see Methods). Across all vessels, anti-PACAP pretreatment reduced the vasodilatory response to PACAP38 during the first 10-min post-injection imaging period compared with vehicle pretreatment (estimated treatment difference = −0.619 z-score units, *P* = 0.0046; Fig. 4a). Anti-PACAP pretreatment did not show evidence of a size-dependent effect (Fig. S3).

**Figure 4.**
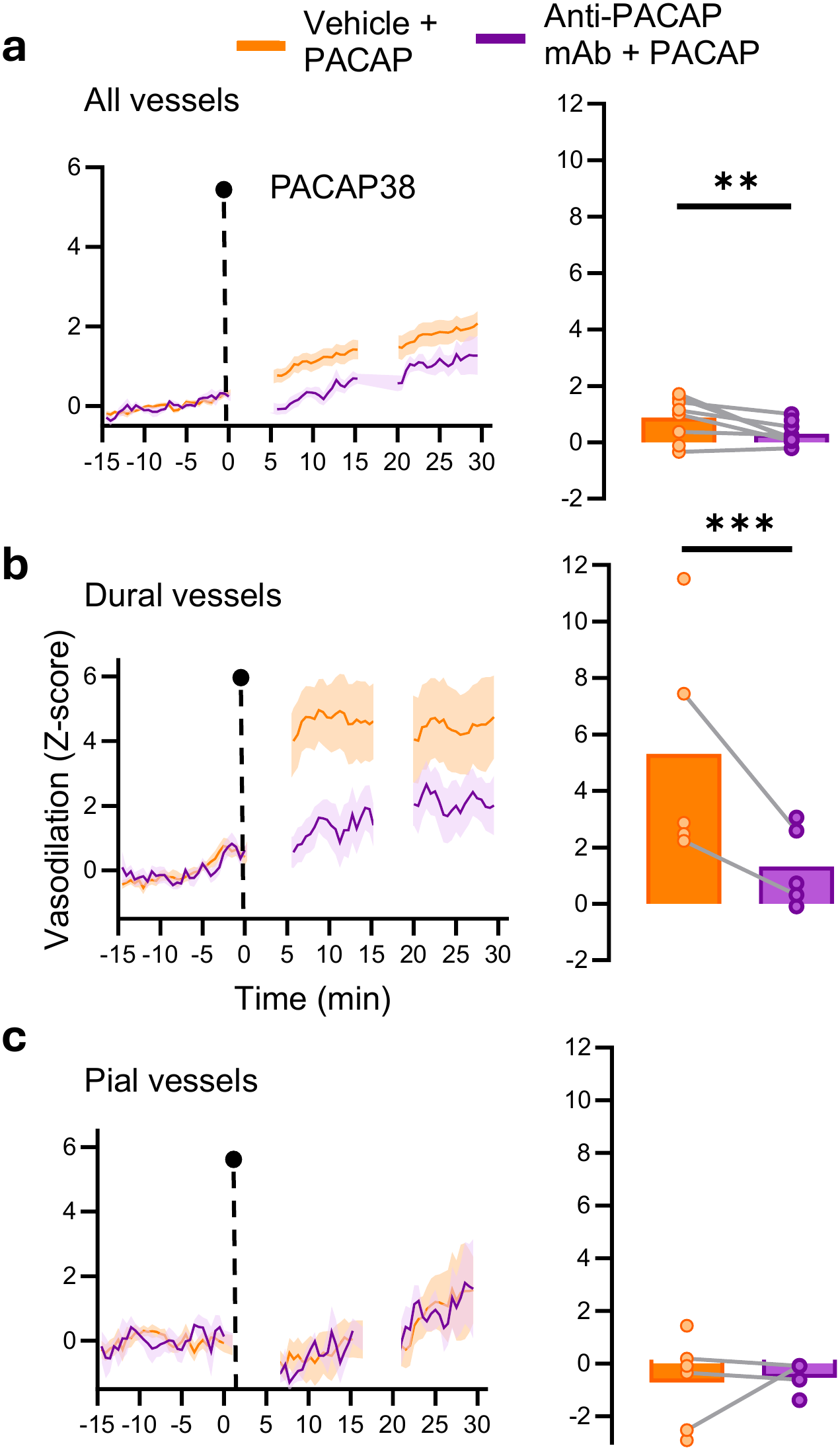
Anti-PACAP monoclonal antibody reduces PACAP38-induced meningeal vasodilation, with a prominent effect in dural vessels. **a,** PACAP38-induced vascular responses across all vessels in mice pretreated with vehicle or anti-PACAP monoclonal antibody. Left, time courses expressed as z-scores relative to the 15-min baseline. The dashed line indicates PACAP38 administration at 0 min, and the gap corresponds to the 5-min pause between imaging periods. Lines show the mean and shaded regions indicate s.e.m. Right, mouse-level mean vasodilation during the first 10-min post-injection imaging period; lines connect measurements from the same mouse. Vehicle + PACAP38, n = 9 mice; anti-PACAP monoclonal antibody (mAb) + PACAP38, n = 9 mice; 7 mice contributed to both conditions. Anti-PACAP pretreatment reduced PACAP38-induced vasodilation (estimate = −0.619 z-score units, *P* = 0.0046). **b,** PACAP38-induced responses in dural vessels, displayed and quantified as in a. Vehicle + PACAP38, n = 5 mice; anti-PACAP mAb + PACAP38, n = 5 mice; 2 mice contributed to both conditions. Anti-PACAP pretreatment reduced PACAP38-induced vasodilation in dural vessels (estimate = −4.054 z-score units, Holm-adjusted *P* = 0.0003). **c,** PACAP38-induced responses in pial vessels, displayed and quantified as in a. Vehicle + PACAP38, n = 6 mice; anti-PACAP mAb + PACAP38, n = 4 mice; 3 mice contributed to both conditions. Anti-PACAP pretreatment did not significantly alter PACAP38-induced vasodilation in pial vessels (estimate = 0.418 z-score units, Holm-adjusted *P* = 0.6881). Partially paired comparisons (a–c) were analyzed using maximum-likelihood linear mixed-effects models with treatment as a fixed effect and mouse identity as a random intercept. Treatment contrasts were evaluated using two-sided Wald tests. Holm correction was applied across the dura and pia compartment-specific contrasts. *P < 0.05; **P < 0.01; ***P < 0.001.

We next examined PACAP neutralization effects separately in dural and pial vessels. In dural vessels, anti-PACAP pretreatment attenuated vasodilation to PACAP38 (estimated treatment difference = −4.054 z-score units, Holm-adjusted P = 0.0003; Fig. 4b). In pial vessels, PACAP38 alone again produced little vasodilation, consistent with the preceding analysis (Fig. 3h), and anti-PACAP pretreatment had no significant effect (estimate = 0.418 z-score units, Holm-adjusted P = 0.6881; Fig. 4c). The treatment × vascular-compartment interaction was significant (Wald χ²(1) = 10.351; P = 0.0013), indicating that the effect of anti-PACAP pretreatment differed between dural and pial vessels. Thus, anti-PACAP pretreatment attenuated the dural vasodilatory response to PACAP38.

## Discussion

High-spatiotemporal-resolution imaging of individual meningeal vessels in freely moving, non-anesthetized mice revealed preferential dural vascular dilation by systemic PACAP38 and showed that this vasodilation can be attenuated by a therapeutically relevant anti-PACAP monoclonal antibody. Although nitroglycerin, levcromakalim, and PACAP38 produced a shared net vasodilatory response, the distribution of responsiveness across individual vessels differed between agents. For PACAP38, anatomical compartment better accounted for the response pattern than vessel caliber alone, identifying the dura as a preferentially PACAP-responsive peripheral meningeal compartment. Demonstrating pharmacological modulation of this dural phenotype provides proof of concept that vessel-resolved vasodilation can report functional PACAP pathway inhibition in vivo, supporting the mechanistic interrogation, drug discovery, and forward and back-translation of PACAP-targeting therapies.

Previous approaches have measured vascular responses to migraine provocateurs, but characterizing these responses in freely moving, non-anesthetized mice allow spatial and temporal dynamics to be examined under conditions that better preserve physiological influences relevant to migraine pathophysiology. The chronic cranial window miniscope preparation allowed the same vessel segments to be tracked during pharmacological challenge without anesthesia or head restraint, while preserving the broader meningeal vascular network and its in vivo physiological context (Degel et al., 2024; Ghosh et al., 2011). This is particularly relevant for clinical pharmacological studies because anesthesia alters vascular tone and neurovascular coupling and can directly affect vasoactive signaling pathways, including KATP-, Ca²⁺-, and NO-dependent mechanisms, which overlap with pathways engaged by the migraine provocateurs examined here (Claassen et al., 2021; Gao et al., 2017; Masamoto & Kanno, 2012; Wen et al., 2026).

Under these conditions, net vasodilation could be resolved into heterogeneous responses across individual vessels within the miniscope field of view, and their distribution examined across dural or pial compartments. Thus, the main advantage of the approach was not simply measurement under more physiological conditions, but conversion of a coarse vascular endpoint into a spatially resolved phenotype suitable for mechanistic and pharmacological interrogation. Mechanistically distinct migraine provocateurs converged on a common net meningeal vasodilatory response, but vessel-level analysis revealed marked, agent-specific heterogeneity in how that response was distributed. This shared net effect parallels human magnetic resonance angiography studies showing extracerebral meningeal arterial dilation across diverse molecular migraine triggers and extends the initial proof-of-concept observations of Degel et al. (2024) across multiple provocateurs (Arquizan et al., 2025).

Individual vessel responses ranged from pronounced dilation to little change or constriction, indicating that similar mean responses can arise from different patterns of vascular recruitment. Changes in meningeal vessel diameter therefore provide a measurable physiological readout of engagement of diverse migraine-relevant pathways, but population averages do not reveal which vascular populations are recruited. This distinction is biologically relevant because restricting analysis a priori to selected vessel classes may overlook responsive populations, whereas averaging across vessels can obscure spatially organized effects. Trigeminal sensory innervation extends beyond major dural arteries to pial vessels and pia-arachnoid tissues, and mechanical stimulation of the pia or small cerebral vessels can evoke referred head pain (Fontaine et al., 2018; Levy & Moskowitz, 2023; Terrier et al., 2021). Broad vessel sampling therefore allowed us to examine whether characteristics of the vessels themselves were associated with differential responsiveness. As an initial stratification, we examined responses by vessel caliber. Nitroglycerin responses tended to be greater in larger vessels, whereas levcromakalim showed no significant size-dependent effect. PACAP38 responses were greater in smaller vessels. As dural vessels were generally smaller than pial vessels, the apparent association between smaller caliber and greater PACAP38 responsiveness could partly reflect anatomical compartment. Examining PACAP38 responses in dura and pia provides a more biologically informative characterization of this pattern than caliber alone.

Systemic PACAP38 preferentially dilated dural over pial vessels, distinguishing it from the broader compartmental responses to nitroglycerin and levcromakalim and supporting a peripheral meningeal component of PACAP action. Nitroglycerin showed no clear compartmental bias, whereas levcromakalim tended toward greater pial responses, but both agents affected vessels in each compartment. This cross-compartmental levcromakalim response is consistent with preclinical dilation of both dural and pial arteries (Gozalov et al., 2005). Human MRA studies also report extracerebral and cerebral dilation after nitroglycerin, whereas cerebral responses to levcromakalim vary across studies (Arquizan et al., 2025). By contrast, PACAP38 preferentially dilated dural vessels, and no clear caliber-response relationship remained within either compartment. PACAP responsiveness is not confined to the dura: PACAP-immunoreactive fibers have been identified around pial and cerebral vessels, topical PACAP38 dilates pial arterioles in vivo, and isolated cerebral arteries respond to PACAP ex vivo (Baun et al., 2011; Grände et al., 2013; Lenti et al., 2007; Lund & Hannibal, 2022; Uddman et al., 1993). Because pial arterioles participate in cerebral autoregulation, changes in systemic blood pressure may contribute to the observed pial responses to these agents. However, this potential contribution could not be assessed because blood pressure was not recorded (Klein et al., 2022). Nevertheless, the limited pial response to systemic PACAP38, despite established pial responsiveness under local and ex vivo conditions, highlights the distinction between the capacity of a vessel to respond to PACAP and whether that response is recruited following systemic exposure in vivo. Mouse tracer studies indicate that PACAP38 can cross the blood–brain barrier after intravenous injection (Banks et al., 1993; Rhea et al., 2018). In the earlier study, approximately 0.053% of the injected radiolabeled dose was detected in whole brain after 5 min (Amin & Schytz, 2018; Banks et al., 1993). Rapid degradation and efflux may nevertheless limit the central availability of intact PACAP38 (Amin & Schytz, 2018). The limited early pial response may therefore reflect insufficient local exposure to PACAP38 or limited responsiveness in vivo; the present experiments cannot distinguish between these possibilities. Our findings demonstrate engagement of the peripheral meningeal vasculature at the dose administered, while later pial responses cannot be excluded. A broadly similar vascular pattern has been observed in human provocation studies, where systemic PACAP38 produces immediate and sustained (>5 hours) prominent dilation of the dural middle meningeal artery with little or no detectable middle cerebral artery dilation (Amin et al., 2012; Amin & Schytz, 2018; Arquizan et al., 2025). Although human MRA assesses larger cerebral vessels than the vessels imaged here, both approaches identify a prominent dural response to systemic PACAP38. Our vessel-resolved imaging extends that large-artery comparison to smaller dural and pial vessels within the imaged field. Complementary electrophysiological evidence shows that systemic PACAP38 activates and sensitizes meningeal Aδ- and C-fiber nociceptors and that local dural lidocaine attenuates ongoing PACAP38-induced activity (Christensen et al., 2025). Because the vascular response emerged within 10 min whereas nociceptor activation developed later, PACAP38 may recruit vascular and sensory components of the peripheral meninges over distinct phases. Together, these observations place preferential early dural vasodilation within a temporally evolving peripheral meningeal response to PACAP38 and provide a rationale for testing whether this vascular response can be attenuated in vivo.

PACAP neutralization provided the key proof of concept that the dural vascular phenotype can report functional PACAP pathway inhibition in vivo. Pretreatment with an anti-PACAP monoclonal antibody attenuated PACAP38-induced meningeal vasodilation, with the treatment effect most evident in dural vessels, where PACAP38-induced dilation was greatest in controls. Thus, the phenotype is not merely descriptive but is pharmacologically modifiable and can serve as a functional readout of PACAP pathway inhibition. Such a readout complements measurements of drug exposure and molecular target binding by testing whether an intervention produces the expected physiological effect in vivo. This concept has direct clinical precedent: PACAP38-induced cranial arterial dilation was used as a prespecified pharmacodynamic endpoint to demonstrate functional PACAP pathway inhibition by Lu AG09222 in humans (Rasmussen et al., 2023), while subsequent phase 2 development demonstrated clinical efficacy of PACAP-targeted therapy in migraine (Ashina et al., 2024). This technology therefore supports a bidirectional translational strategy: candidate therapeutics can be tested prospectively for their ability to suppress a defined PACAP-responsive phenotype, while clinically observed pharmacodynamic responses can be modelled and dissected at vessel-level resolution in preclinical studies. Suppression of vasodilation should not be considered a surrogate for clinical efficacy; rather, it provides a functional pharmacodynamic readout of PACAP pathway inhibition that may bridge therapeutic intervention and clinical outcome. By linking a clinically relevant PACAP-responsive phenotype to vessel-specific biology and therapeutic modulation, this approach provides a framework for functional pathway interrogation, drug discovery, and the forward and back-translation of migraine therapeutics.

### Limitations of the study

Vascular responses were measured without concurrent migraine-relevant behavioral outcomes; attenuation of PACAP38-induced dural vasodilation should therefore be interpreted as a functional pharmacodynamic readout of PACAP pathway inhibition, not as evidence that vasodilation mediates migraine pain or predicts clinical efficacy. Locomotion and arousal may contribute to vessel-diameter variability. In head-fixed awake mice, locomotion has been associated with dural constriction and pial dilation (Gao & Drew, 2016). Locomotor state was not included in the primary analyses, although study-matched vehicle controls help account for this potential source of variability. Blood pressure was not recorded, limiting assessment of systemic hemodynamic contributions to the observed vascular responses.

Systemic PACAP38 also bypasses endogenous peptide release during spontaneous migraine. Nevertheless, provocation models provide controlled systems for forward and back-translation (Ashina et al., 2022). Dural and pial vessels were manually classified using conservative morphological criteria, limiting compartment-specific analyses to confidently classified vessels while reducing the risk of misclassification. Arterial and venous identity was not systematically resolved.

Generalizability across sex, species, and age remains to be determined.

## Resource Availability

### Materials availability

The anti-PACAP tool monoclonal antibody used in this study was provided by H. Lundbeck A/S. Enquiries regarding access should be directed to the lead contact and are subject to approval by H. Lundbeck A/S.

### Data and code availability

- The authors do not have permission to share data.

- All original code has been deposited at https://github.com/LEWN-HLU/meningeal-vessel-diameter-analysis.git

- Any additional information required to reanalyze the data reported in this paper is available from the lead contact upon request.

## Acknowledgments

This work was supported by H. Lundbeck A/S.

## Author Contributions

Conceptualization: P.B., C.D, B.J.H., A.A.; Methodology: L.S., K.Z., J.Z, D.O., J.N.; Software: L.S., Validation: L.S., C.D.; Formal analysis: L.S.; Investigation: C.D., T.B., K.Z., F.W.; Data curation: L.S.; Resources: B.J.H.; Visualization: L.S.; Writing: L.S, P.B.; Supervision: P.B., S.L.C.

## Methods

### EXPERIMENTAL MODEL AND SUBJECT DETAILS

#### Animals

This study presents a secondary analysis of data from a previously established cohort of male mice. The sample size was determined by the number of eligible animals in the existing dataset; no a priori power calculation was performed. No additional animals were used for the present analyses, consistent with the Reduction principle of the 3Rs. Because the cohort included only male mice, this study was not designed to assess sex-dependent vascular responses.

#### Experiments conducted at Inscopix

Male C57BL/6JCrl mice aged 12–14 weeks at the time of surgery were used for experiments conducted at Inscopix (Mountain View, CA, USA). Animals were group housed before surgery and pair housed after surgery using cage dividers that permitted social interaction while preventing access to implanted headcaps and microscope baseplates. Mice were maintained on a reversed 12-h light–dark cycle, with lights off from 06:00 to 18:00. All procedures were conducted under protocols approved by the Institutional Animal Care and Use Committee of LifeSource Biomedical Services, LLC under protocol INS-22-002. Data produced by Inscopix are shown in Fig. 2c and supplementary Fig. 4a, the rest was collected at Lundbeck.

#### Experiments conducted at Lundbeck

Male C57BL/6JCrl mice aged 6-8 weeks and weighing 20–30 g were obtained from Charles River (Germany). Mice were housed individually throughout the study in Macrolon type III low cages (20 x 35 x 18 cm) containing nesting material and standard environmental enrichment, with food and water available ad libitum. Animals were acclimatized for at least 5 days, often up to 2 weeks, before experimental procedures. Mice were maintained on a 12-h light–dark cycle, with lights on at 06:00. All procedures were approved by the Danish Animal Experiments Inspectorate (Dyreforsøgstilsynet) under permit 2025-04513 and were conducted in accordance with Danish legislation on animal experimentation and Directive 2010/63/EU.

## METHOD DETAILS

### Preparation and administration of vascular contrast agent

For imaging experiments conducted at both sites, fluorescein isothiocyanate–dextran (2,000 kDa; Fluorescein isothiocyanate–dextran, Sigma-Aldrich, Missouri, USA, #FD2000S) was dissolved in sterile 0.9% NaCl at a final concentration of 9 mg/ml. The solution was passed through a sterile 5 µm nylon syringe filter (NYS502550, Sterlitech, Auburn, WA, USA), protected from light, and stored at 4°C for the duration of the acute study. During imaging sessions, the solution was maintained on ice until administration.

Before imaging, mice were warmed for approximately 3-5 min to promote tail-vein dilation and were weighed. FITC-dextran was administered through the lateral tail vein using a 28-G insulin syringe (New Jersey, NY, USA) at a volume of approximately 150-200 µl 15 min before baseline imaging. Successful intravenous administration was confirmed by blanching of the tail vein during injection and subsequent visualization of fluorescent meningeal vessels.

### Chronic cranial window surgery and miniscope baseplate implantation

#### Procedures conducted at Inscopix

Mice were anesthetized with isoflurane in oxygen (4–5% induction, 1–2% maintenance) and secured in a stereotaxic frame (David Kopf Instruments, Tujunga, CA, USA) on a heating pad maintained at 37°C. Ketofen (5 mg/kg, subcutaneous) and carprofen (2.5 mg/kg, subcutaneous) were administered before surgery. Following a midline scalp incision, the skull was exposed and cleaned.

A craniotomy slightly larger than 3 mm in diameter was made in the right parietal bone adjacent to the sagittal and coronal sutures using a K.1070 High Speed Rotary Micromotor Drill (Foredom, Danbury, CT, USA) fitted with a 2.1-mm carbon steel burr (19007-21, Fine Science Tools, Foster City, CA, USA). The dura mater was preserved throughout the procedure. A 3-mm no. 0 round glass coverslip (CS-3R-0; Warner Instruments, Harvard Bioscience, Holliston, MA, USA) was positioned over the craniotomy and secured with cyanoacrylate adhesive (Loctite Super Glue Gel Control, SKU 234790; Henkel Corporation, Westlake, OH, USA).

For miniscope alignment and baseplate implantation, an nVista 3.0 miniature microscope (Inscopix, Mountain View, CA, USA) modified to provide a working-distance range of 200–500 µm from the objective lens was mounted on the stereotaxic arm. A compatible baseplate was attached to the microscope, and the live image stream was monitored using Inscopix Data Acquisition Software (IDAS, Inscopix, Palo Alto, CA, USA). During alignment, images were acquired at 10 Hz using an excitation LED power of 1.5 mW and a gain setting of 8 under ambient overhead. The miniscope position and angle were adjusted until the objective lens was parallel to the coverslip, and meningeal vessels were visualized near the midpoint of the working-distance range. The baseplate was then fixed to the skull using C&B Metabond® Quick Adhesive Cement System (Parkell, Inc., Edgewood, NY, USA).

After curing, the microscope was removed and a protective baseplate cover was installed.

Following surgery, mice recovered in a warmed cage and were monitored until ambulatory. Animals were monitored daily for at least 3 days. Imaging experiments began 3–4 weeks after surgery. Animals were excluded if inflammation or infection was observed at the cranial window site, including tissue opacity or clouding, or if meningeal vessels could not be visualized reliably.

#### Procedures conducted at Lundbeck

FITC-dextran was administered intravenously before surgery to visualize the meningeal vasculature and guide miniscope field-of-view selection and baseplate placement. The surgical field was prepared using aseptic procedures. Anesthesia was induced with 5% isoflurane delivered with oxygen and nitrous oxide at flow rates of 0.3 and 0.7 l/min, respectively, and maintained at approximately 2%. Mice were secured in a stereotaxic frame (David Kopf Instruments, Tujunga, CA, USA) on a heated pad set to maintain body temperature at 37°C. Ophthalmic ointment (Actavis, Søborg, Denmark) was applied, and Marcain (2.5 mg/ml bupivacaine, AstraZeneca, Albertslund, Denmark) was administered subcutaneously beneath the planned incision site. Noromox (150 mg/ml amoxicillin hydrate, ScanVet, Fredensborg, Denmark), and Norodyl (50 mg/ml carprofen, ScanVet, Fredensborg, Denmark) were administered subcutaneously. Depilatory cream (Veet Pure Hair Removal Cream, Normal Skin, Body & Legs; Reckitt, Søborg, Denmark) was applied to the surgical area and left for two minutes, then removed with cotton tips. The exposed skin was disinfected with ethanol swabs.

A midline scalp incision was made, and the subcutaneous tissue was gently retracted using sterile cotton tips to expose the skull. The skin margins were secured to the skull using cyanoacrylate adhesive (Loctite Super Glue Power Gel, 3 g; EAN 5010266241159; Henkel Denmark A/S, Ballerup, Denmark). A 4-mm cranial window was positioned on top of the skull over the right somatosensory cortex lateral to bregma and outlined with a semi-permanent marker. The skull surface surrounding the intended craniotomy was gently scored with a sterile scalpel (No. 23; 1902.503; Hounisen Laboratorieudstyr A/S, Denmark) to improve subsequent adhesion.

The outlined skull region was thinned using a surgical drill under sterile saline irrigation, and the central bone flap was removed while preserving the underlying dura mater. Bone dust and any minor bleeding were consistently cleared using sterile saline irrigation, with excess fluid absorbed using sterile cotton tips. The exposed dura was maintained under sterile saline. Animals were excluded and euthanized if dural damage or substantial hemorrhage occurred.

A 4-mm cover glass was positioned directly over the dura and secured with superglue to the skull bone. For miniscope alignment and baseplate implantation, an nVue2.0 miniscope (Inscopix, Palo Alto, CA, USA) attached to its baseplate (Inscopix, Palo Alto, CA, USA) was mounted on the stereotaxic arm. The microscope position, angle, focal plane and magnification were adjusted until meningeal vessels were clearly visualized. The baseplate was fixed to the skull using RelyX Unicem Aplicap dental cement (RelyX Unicem Aplicap, 3M, Saint Paul, MN, USA). After curing, the microscope was removed and a protective cap (Inscopix, Palo Alto, CA, USA) was secured with a small screw.

Following surgery, mice recovered in a clean cage on a heating pad maintained at 37°C and were provided with moistened food and one-quarter of a Rimadyl MD tablet (2 mg/Tablet Rimadyl, Bio-Serv, Fleming ton, New Jersey, USA). Motor recovery and body weight were monitored postoperatively. Amoxicillin was administered subcutaneously for 5 days after surgery at 1 ml/kg and carprofen at 20 mg/kg if the Rimadyl MD tablet was not eaten. Animals were excluded if inflammation or infection was observed at the cranial window site, including tissue opacity or clouding, or if meningeal vessels could not be visualized reliably.

### Habituation

#### Experiments conducted at Inscopix

To habituate animals to experimental procedures, mice underwent habituation sessions on two consecutive days during the week prior to imaging. Each habituation session consisted of 5 min of restraint in the Tailveiner apparatus, 60 min exploration of the 42 x 42 cm square open field arena while attached to the miniscope, and intraperitoneal injection of DPBS (10 mL/kg).

#### Experiments conducted at Lundbeck

Mice underwent habituation for 1 h per day over 5 consecutive days. During each session, mice were gently handled and placed in an assigned Macrolon type III low cage while carrying a dummy miniscope (Inscopix, Palo Alto, CA, USA) attached to the baseplate. The same cage was used throughout habituation and subsequent experiments. Animals reused in later experiments underwent one additional day of re-habituation with the dummy miniscope in the same cage before imaging.

### Miniscope imaging of meningeal vasculature

#### Experiments conducted at Inscopix

On imaging days, mice received tail-vein administration of FITC-dextran as described above. Mice were briefly restrained while the protective baseplate cover was removed and the nVista3.0 miniscope (Inscopix, Palo Alto, CA, USA) was docked onto the implanted baseplate, after which they were transferred directly to the open-field arena.

Imaging parameters were adjusted for each session using the live image histogram in IDAS (Inscopix, Palo Alto, CA, USA) to maximize fluorescence signal while avoiding pixel saturation (optimal range: 90-95%). Recordings were acquired at 10 Hz using an excitation LED power of 0.1–0.4 mW and gain values of 1.2–5.3.

#### Experiments conducted at Lundbeck

Imaging was performed more than 4 weeks after cranial-window implantation. On imaging days, mice received tail-vein administration of FITC-dextran as described above. Meningeal vessels were imaged using an nVoke 2.0 or nVue miniature microscope (Inscopix, Palo Alto, CA, USA) mounted on the implanted baseplate, with a commutator used to minimize cable entanglement. See Figure 1a and supplementary Fig. S1 for schematic illustrating experimental paradigms. The experiments were conducted within a sound-proof chamber, with lighting conditions set to 26 LUX.

The field of view was selected using IDAS (Inscopix, Palo Alto, CA, USA) and maintained for each mouse across imaging sessions. Recordings were acquired at 10 Hz using an excitation LED power of 0.3–1.2 mW and gain values of 1–3. Excitation power and gain were adjusted to maximize fluorescence signal while avoiding pixel saturation, with the live-image histogram maintained within an optimal range of 60–70%. FITC-dextran fluorescence was imaged using the green imaging channel of nVue or nVoke miniature miniscopes. Imaging recordings were acquired in discrete blocks rather than continuously, with 5-minute intervals between blocks. One 15-minute block was recorded at baseline, followed by up to three 10-minute blocks to monitor vascular responses after agent administration. The intervals between blocks allowed verification of the focal plane while minimizing the risk of fluorescence photobleaching caused by continuous miniscope LED illumination.

### Experimental design

#### Experiments conducted at Inscopix

For the PACAP dose-escalation study, each mouse received sequential PACAP38 doses of 0.3, 0.6, and 1.2 mg/kg within the same dose-escalation experiment, resulting in a repeated-measures design.

PACAP38 (0.3, 0.6 and 1.2 mg/kg, Bachem, Cat. No. 4031157), and its vehicle were administered at 10 ml/kg. PACAP38 was prepared in DPBS. All compounds were administered intraperitoneally. Compounds were prepared fresh on the day of imaging, protected from light, and maintained on ice until administration. Each imaging session consisted of a 15-min baseline recording followed by compound administration and up to 45 min of post-injection imaging. Imaging sessions were separated by 6–9 days.

#### Experiments conducted at Lundbeck

Levcromakalim (1 mg/kg, i.p.) and nitroglycerin (10 mg/kg, i.p.) were selected based on their use in mouse migraine-provocation studies (Bates et al., 2010; Christensen et al., 2021; Guo et al., 2022; Pradhan et al., 2014). For PACAP38, published mouse studies use different systemic doses and routes for different outcomes, including 0.3 mg/kg i.p. for light aversion and 2–30 µg/kg s.c. for tactile hypersensitivity (Ernstsen et al., 2022; Kuburas et al., 2021). The initial Inscopix study (Supplementary Fig. 4a) served as an exploratory screen of PACAP38-induced vascular responses across increasing doses (0.3, 0.6, and 1.2 mg/kg). Responses were variable at 0.3 mg/kg and appeared more pronounced at higher doses. Because the same mice received the doses sequentially, the study did not isolate the response to a single acute dose. We therefore tested lower doses in-house under the acute-challenge conditions used here (Supplementary Fig. 4b). Both 0.1 and 0.3 mg/kg produced significantly greater vasodilatory responses than vehicle, with no statistically detectable difference between them; 0.1 mg/kg was selected as the lowest tested dose producing a significant response.

Study designs differed across pharmacological experiments. The levcromakalim (MedChemExpress, New Jersey, USA #HY-14255) study used independent treatment and vehicle groups. The nitroglycerin (Cambrex, Wiesbaden, Germany) study used a fully paired design in which the same mice received nitroglycerin and vehicle in separate imaging sessions; this study was performed blinded to treatment condition, whereas the other in-house pharmacological studies were not blinded. For PACAP38 (Bachem, CA, USA #4031157), the 0.1 mg/kg comparison used independent groups and the PACAP38 versus anti-PACAP monoclonal antibody study were partially paired, with some mice contributing to both conditions and others to only one. For the PACAP38 compartment analysis, PACAP38-treated observations from the acute-dosing and anti-PACAP studies were pooled. Animals reused in subsequent experiments underwent an additional day of re-habituation with the dummy miniscope before imaging, with a washout period of at least 7 days between experiments. FITC-dextran was administered intravenously before imaging as described above. Levcromakalim (1 mg/kg) was prepared in 0.9% saline, nitroglycerin (NTG/GTN; 10 mg/kg) in 12.2% ethanol in saline, and PACAP38 (0.1 mg/kg) in 0.9% saline. All compounds were administered intraperitoneally. Compounds were prepared fresh on the day of imaging, protected from light, and maintained on ice until administration. Following a 15-min baseline recording, the pharmacological challenge was administered and meningeal vasculature was imaged. Post-injection imaging included up to three 10-min recording periods separated by 5-min pauses, during which the miniscope focal plane could be realigned if required. For anti-PACAP experiments, anti-PACAP monoclonal antibody (30 mg/kg, i.p.; 10 mL/kg) was administered 24 h before the PACAP38 challenge. The same antibody and pretreatment schedule were used in a previous mouse PACAP38 provocation study (Guo et al., 2022).

### Histology and tissue processing

#### Whole-head clearing and light-sheet fluorescence microscopy

Whole-head tissue clearing and light-sheet fluorescence microscopy were performed by Gubra (Hørsholm, Denmark). Male C57BL/6JRj mice were transcardially perfused with heparinized PBS followed by 10% neutral buffered formalin (NBF). Heads were decapitated, skinned, and the jaw and surrounding muscle tissue were removed, then post-fixed overnight in 10% NBF, washed, and stored at 4°C in PBS/KPBS containing 0.02% sodium azide. Samples used in Figure 3a were decalcified for 7 days in 20% EDTA with daily solution changes and processed using a modified iDISCO protocol (Perens et al., 2021), including methanol dehydration, dichloromethane delipidation, bleaching, permeabilization, collagenase digestion, blocking, and incubation with directly conjugated antibodies against CD31, von Willebrand factor (vWF), and SM22α, followed by clearing in ethyl cinnamate. Cleared heads were imaged using an LCS SPIM light-sheet microscope with a 4× C objective, and three-dimensional visualization was performed using Imaris or Aivia software.

#### Euthanasia and tissue collection

Mice used for histological analyses were euthanized and transcardially perfused. Animals first received an intraperitoneal overdose of Avertin (approximately 0.4 ml per mouse). The heart was then perfused for 2 min with 24 ml KPBS (pH 7.4), which for selected animals contained 0.015 mg/ml lectin-binding dextran: Wheat Germ Agglutinin (WGA) (Invitrogen, Alexa Fluor 488 or 647), followed by 10% neutral buffered formalin (NBF) for 2 min.

Following fixation, mice were decapitated, and the skull cap (calvaria) with the attached dura mater was dissected using a previously described method (Nilsson et al., 2022). Brains were immersion-fixed overnight in NBF at room temperature, washed three times for 10 min in KPBS on a horizontal shaker, and stored at 4°C in KPBS containing 0.02% sodium azide until further processing.

#### Dura mater dissection

The dura mater was dissected from the calvaria using fine forceps under a Leica S9i Digital Stereo Microscope. The tissue was gently loosened around the perimeter of the calvaria, removed, transferred to 24-well plates containing 500 µl KPBS, and washed three times for 10 min before immunostaining.

#### Immunohistochemistry

Dura mater samples were washed three times for 10 min in KPBS and blocked for 1 h at room temperature in 300 µl blocking buffer containing 5% normal donkey serum. Samples were then incubated overnight at 4°C on a horizontal shaker with anti-CD31 antibody (1:100, rat, Invitrogen, MA1-40074).

The following day, samples were washed three times for 10 min in KPBS containing 0.25% BSA and 0.1% Triton X and incubated for 1 h at room temperature with the corresponding secondary antibody: anti-rat Alexa Fluor 568 (1:1000, Invitrogen, A-11077). Samples were then washed three times for 5 min in KPBS.

The dura was then mounted on Superfrost Plus slides (Süsse, Gudensberg, Germany) and coverslipped with ProLongTM Diamond Antifade Mountant with DAPI (Thermo Fisher Scientific, Waltham, Massachusetts, USA).

All immunofluorescence imaging was acquired using Leica Mica Microhub Imaging System.

#### Miniscope histological imaging

Dissected brains and dura from animals perfused with WGA (Invitrogen, Alexa Fluor 488) were positioned beneath an nVue miniscope mounted on a stereotaxic arm. The uppermost pial layer of the brains and, separately, dura were imaged using the green fluorescence channel, to visualize pial blood vessels on the cortical surface and dural blood vessels (Fig. 3c). Recordings were acquired at 10 Hz using an excitation LED power of 0.3–1.2 mW and gain values of 1–3.

## QUANTIFICATION AND STATISTICAL ANALYSIS

### Data processing and vessel diameter estimation

Raw imaging data were processed using Inscopix Data Processing Software (Inscopix, Mountain View, CA, USA). One-photon imaging videos of baseline and post-drug recordings from each session were concatenated and analyzed as a continuous time series. Pre-processing comprised band-pass filtering and motion correction of the raw imaging video, followed by vessel diameter annotation and subsequent post-processing analysis.

### Pre-processing

#### Image filtering and motion correction

To correct for defective pixels and remove artefacts, a 3x3 median filter was applied to the recordings. They were then subjected to spatial filtering to improve image quality and visual clarity. The spatial band-pass filter was specified in oscillations per pixel rather than in physical distance. No low-frequency cut-off was applied, whereas the high-frequency cut-off was set to 0.095 pixel⁻¹. Filtering was performed independently on each frame by subtracting a more strongly smoothed version of the image from a less strongly smoothed version, as described in the Inscopix Data Processing Software User Guide (IDPS, Release 1.9.5, Inscopix, Palo Alto, CA, USA).

Finally, motion correction was applied to compensate for frame-to-frame displacement and maintain consistent spatial correspondence throughout each recording. Typically, the complete first frame served as the reference region. For recordings with pronounced movement within the field of view, a mean image generated from the full recording was used as a reference to improve the accuracy of the correction.

### Vessel diameter estimation

Following motion correction, vessels were manually annotated in IDPS using lines drawn perpendicular to individual vessel segments, with both endpoints positioned outside the vessel boundaries. These lines defined the pixel-intensity profiles used for vessel edge detection and diameter estimation. Annotation lines were included only if they intersected a single vessel throughout the recording. One annotation was generated per vessel segment, while downstream branches were treated as separate vessel segments. Where possible, annotation endpoints were positioned in regions that remained substantially darker than the vessel lumen throughout the recording to improve edge detection.

Vessel diameter was estimated using methods described in Degel et al. (2024), with a custom analysis algorithm (Inscopix, Palo Alto, CA, USA). For each annotated line and imaging frame, the pixel-intensity profile was used to identify local minima on either side of the vessel lumen and the maximum global intensity within the profile. Vessel edges were defined as the pixels nearest the half-maximum intensity between the global maximum and the corresponding local minimum on each side of the vessel. Diameter was calculated as the distance between the two detected vessel edges. Diameter estimates were visually inspected for consistency with the apparent vessel boundaries.

Diameter was estimated for every acquired frame at 10 Hz and subsequently averaged across consecutive 1-s intervals (10 frames) to generate diameter traces at 1 Hz temporal resolution.

### Post-processing

Following the diameter estimation in IDPS, a csv file was extracted for each animal/session and used for further data processing. Post-processing and visualization were performed using custom Python scripts (Python version 3.12.13) and GraphPad Prism (version 10.1.2, GraphPad Software, San Diego, CA, USA). The post-processing script was developed with assistance from OpenAI Codex. If not specified, data are shown as mean ± standard error of the mean (s.e.m.).

### Vessel diameter normalization and response quantification

For each accepted vessel, the 1-Hz diameter trace from the csv file was normalized to that vessel’s own 15-min baseline period. The baseline mean and standard deviation were calculated from this period, and each diameter measurement was converted to a z-score according to

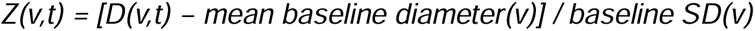

where *D(v,t)* is the diameter of vessel *v* at time *t*, and *mean baseline diameter(v)* and *baseline SD(v)* are the mean and standard deviation of that vessel’s baseline diameter, respectively. This generated a time-resolved z-score trace for each vessel.

Thresholds were defined to classify vasoconstrictive and vasodilatory responses. For each vessel, the z-score quantified the deviation of the response from its baseline mean in standard-deviation units. Responses with mean z-scores ≤ −1.96 or ≥ 1.96 were classified as vasoconstrictive or vasodilatory, respectively. Negative and positive z-scores indicated the direction of diameter change; values between these thresholds did not meet either response criterion.

For each 10-min response period following the baseline, z-scores were averaged across time to generate one response value per vessel. Vessel-level response values were subsequently averaged within each animal/session to generate the value used for statistical analysis, where applicable.

As an additional outcome measure, percent change was calculated as 100 × (mean diameter during the first post-injection period − mean baseline diameter) / mean baseline diameter. Vessel-level percent changes were then averaged within each animal/session and vessel category to obtain animal-level summary values. Although less intuitive than percent change, the z-score expresses each vessel’s evoked diameter change relative to its spontaneous baseline fluctuations, thereby normalizing differences in baseline variability between vessels; it is therefore used throughout this paper.

Vessels with substantial missing or unreliable diameter estimates were excluded during preprocessing. For otherwise accepted vessels, on postprocessing, isolated missing time points were omitted from calculations of baseline statistics and response averages. If the baseline standard deviation was zero or invalid, z-scores for that vessel were treated as missing.

### Classification into vessel-size classes

To distinguish vessels at the extremes of the observed size distribution, vessels were classified as small or large based on their measured baseline diameter. As shown in Fig. 2b, the diameter distribution was visually right-skewed, with most vessels measuring less than 20 µm. Small vessels were therefore defined as those with diameters <20 µm, whereas large vessels were defined as those with diameters >50 µm. These cut-offs were selected to maximize separation between the two size classes. Vessels with diameters between 20 and 50 µm were considered intermediate-sized and were excluded from this binary classification.

### Morphological classification of dural and pial vessels

To distinguish dural and pial vessels within the meningeal field of view acquired using the Inscopix miniscope, vessels were classified according to predefined morphological criteria. Dural vessels were identified as relatively superficial, usually smaller-caliber vessels with long, comparatively straight trajectories that often formed small loops. Pial vessels were characterized by larger caliber, more frequent branching over relatively short segments, and apparent penetration into deeper tissue layers, inferred from the abrupt termination of the vessel within the imaging plane.

As illustrated in Fig. 3g, this classification strategy was deliberately conservative. Most vessels were therefore not assigned to a specific meningeal layer when their morphology did not provide sufficient evidence for a reliable classification. The large proportion of unclassified vessels provides an indication of the stringency of the approach and limits the risk of misclassifying vessels based solely on ambiguous morphological features.

To aid visual interpretation of dural and pial morphology, an illustrative schematic was created using BioRender’s AI-assisted image-generation tool. The schematic depicts the representative morphological features used to distinguish between the two vessel classes. A reference image representing each meningeal layer (Fig. 3b-c) was first provided to ChatGPT, which was instructed to describe the visible morphological features, and provide instructions for generating an illustrative representation based on those features. These instructions were subsequently refined by incorporating the descriptions of dura and pia from above, the average sizes and range of vessel diameter in each layer (Fig. 3e-f), the relevant anatomical labels and visual specifications, including the assignment of features to the dural or pial layer, the intended color scheme, and the desired visual overlays (XY- and Z-overlay). The final prompt was entered into BioRender’s AI-assisted image-generation tool to create the schematic shown in Fig. 3d. The resulting schematic is intended as an illustrative aid and does not represent a direct reconstruction of the vessels observed in the recordings.

### Statistical analysis

Unless otherwise stated, vessel-level responses were averaged within each animal/session before inferential analysis, and vessels from the same animal were not treated as independent biological replicates. Time-course plots were used to visualize response dynamics and were not tested at individual time points. For analyses testing whether a normalized vascular response differed from baseline, animal-level mean responses over the predefined response period were compared with zero using a two-sided one-sample *t*-test. For PACAP38 responses in dural and pial vessels, these two baseline comparisons were Holm-adjusted together. Statistical analyses were selected according to the dependency structure of each study. Fully paired data sets of migraine inducer versus vehicle were analyzed using a two-sided Wilcoxon matched-pairs signed-rank test. For independent data set comparisons, a two-sided Mann-Whitney U test was used. A linear mixed-effects model was used when data was partially paired or clustered within mice (i.e. comparing vessels from the same mice as with dura and pia), allowing us to retain animals that contributed to only one condition without incorrectly assuming that repeated measurements from the same mouse were independent. To assess the relationship between baseline vessel diameter and vascular response, z-score response was modeled as a function of baseline vessel diameter with mouse identity included as a random intercept; slope *P* values were Holm-adjusted across levcromakalim, nitroglycerin, and PACAP38. Models were fitted by maximum likelihood using MixedLM in statsmodels 0.14.6 under Python 3.12.13, and contrasts were evaluated using two-sided Wald tests. For size-stratified z-score analyses, mixed-effects models included treatment, vessel-size category and their interaction as fixed effects, with mouse identity as a random intercept. Holm correction was applied across the two small- and large-vessel treatment contrasts within each agent, and across the dura- and pia-specific treatment contrasts in the anti-PACAP analysis. Analyses and visualization was performed using custom Python scripts and GraphPad Prism 10.1.2. Statistical significance was defined as *P* < 0.05.

## WORKING KEY RESOURCES TABLE INVENTORY

This is not updated yet and is also not required to be done before acceptance by journal.

## Supplementary

**Supplementary Table 1 Meningeal vascular responses expressed as z-scores and percent change from baseline.** Values are mean ± s.e.m.; n denotes mice. Vessel-level responses during the first 10-min post-injection imaging period were averaged within each animal/session and vessel category. Values are descriptive group summaries, rather than model-estimated treatment differences.

**Supplementary Figure 1 Experimental timelines for meningeal vascular imaging and pharmacological challenge paradigms. a,** Experimental timeline for vasoactive compound challenge experiments. Mice underwent cranial-window surgery and baseplate implantation 4 weeks before imaging. On the imaging day, FITC–dextran was administered intravenously 30 min before compound administration. A 15-min baseline imaging period was followed by intraperitoneal injection of vasoactive compound or vehicle at 0 min and three 10-min imaging sessions separated by 5-min intervals. Levcromakalim, nitroglycerin, and PACAP38 were administered at 1 mg/kg, 10 mg/kg, and 0.1 mg/kg, respectively. **b,** Experimental timeline for anti-PACAP mAb pretreatment experiments. Mice underwent cranial-window surgery and baseplate implantation 4 weeks before imaging. Anti-PACAP mAb or vehicle was administered intraperitoneally 1 day before imaging. On the imaging day, FITC–dextran was administered intravenously 30 min before PACAP38 or vehicle administration. A 15-min baseline imaging period was followed by intraperitoneal injection of PACAP38 or vehicle at 0 min and two 10-min imaging sessions separated by 5-min intervals. PACAP38 was administered at 0.1 mg/kg; anti-PACAP mAb was administered at 30 mg/kg. Created with BioRender.com.

**Supplementary Figure 2 Dural and pial vascular responses to nitroglycerin and levcromakalim.** Time courses of vasodilation in dural and pial vessels following administration of nitroglycerin (left) or levcromakalim (right). Responses are expressed as z-scores relative to the 15-min baseline. Vessel-level responses were averaged within each animal/session and vascular compartment at each time point and then across animals. Lines show the group mean and shaded regions indicate s.e.m. Compound administration occurred at 0 min. Nitroglycerin: dura, n = 2 mice/2 sessions/40 vessels; pia, n = 3 mice/3 sessions/36 vessels. Levcromakalim: dura, n = 4 mice/4 sessions/33 vessels; pia, n = 3 mice/3 sessions/21 vessels. Gaps indicate intervals between imaging sessions. No inferential statistical tests were applied to individual time points.

**Supplementary Figure 3 Time courses and mean vasodilation stratified by vessel size. a,** Small vessels (<20 µm). **b**, Large vessels (>50 µm). Left, time courses expressed as z-scores relative to the 15-min baseline. The dashed line indicates PACAP38 administration at 0 min, and the gap marks the 5-min pause between imaging periods. Lines show the mean and shaded regions indicate s.e.m. Right, mouse-level mean vasodilation during the first 10-min post-injection imaging period; lines connect measurements from the same mouse. In a, vehicle + PACAP38, *n* = 9 mice, and anti-PACAP monoclonal antibody (mAb) + PACAP38, *n* = 9 mice; 7 mice contributed to both conditions. The treatment estimate (anti-PACAP mAb + PACAP38 minus PACAP38) was −0.767 z-score units (s.e. = 0.500, *z* = −1.535, Holm-adjusted *P* = 0.249). In b, vehicle + PACAP38, *n* = 7 mice, and anti-PACAP mAb + PACAP38, *n* = 5 mice; 4 mice contributed to both conditions. The treatment estimate was −0.499 z-score units (s.e. = 0.632, *z* = −0.791, Holm-adjusted *P* = 0.429). The treatment × vessel-size interaction was not significant (Wald χ²(1) = 0.111, *P* = 0.739). Partially paired comparisons were analyzed using maximum-likelihood linear mixed-effects models with mouse identity as a random intercept. Planned treatment contrasts were evaluated using two-sided Wald tests and Holm-adjusted across the two vessel-size categories.

**Supplementary Figure 4 PACAP38 dosing studies show increased meningeal vasodilation. a,** Vascular responses during the Inscopix dosing study, shown as vasodilation z-scores across continuous PACAP38 dosing at 0, 0.3, 0.6 and 1.2 mg/kg. Lines connect measurements from the same mouse (*n* = 8 mice). No statistical test was performed. **b,** Vascular responses following Lundbeck dosing with vehicle or PACAP38 (0.1 or 0.3 mg/kg). Bars show group means and error bars indicate s.e.m. Vehicle, *n* = 16; PACAP38 (0.1 mg/kg), *n* = 11; PACAP38 (0.3 mg/kg), *n* = 6. The overall difference among groups was significant (Kruskal–Wallis test, *P* = 0.0043). Dunn’s multiple-comparisons test showed differences between vehicle and PACAP38 at 0.1 mg/kg (adjusted *P* = 0.0192) and 0.3 mg/kg (adjusted *P* = 0.0228); the two PACAP38 doses did not differ significantly (adjusted *P* > 0.9999). *P* < 0.05.

## Declaration of generative AI and AI-assisted technologies in the writing process

During preparation of this work, the authors used ChatGPT (OpenAI) to revise, proofread, and improve the readability of the manuscript. OpenAI Codex was used to assist with developing the post-processing script. BioRender AI was used to generate the illustrative schematic in Figure 3D. The authors reviewed and edited the AI-assisted text and figure, checked the code and analyses, and take full responsibility for the content of the publication.

## References

Agarwal, N., & Carare, R. O. (2021). Cerebral Vessels: An Overview of Anatomy, Physiology, and Role in the Drainage of Fluids and Solutes. Frontiers in Neurology, 11. 10.3389/fneur.2020.611485

Al-Karagholi, M. A.-M., Hansen, J. M., Guo, S., Olesen, J., & Ashina, M. (2019). Opening of ATP-sensitive potassium channels causes migraine attacks: a new target for the treatment of migraine. Brain, 142(9), 2644–2654. 10.1093/brain/awz199

Amin, F. M., Asghar, M. S., Guo, S., Hougaard, A., Hansen, A. E., Schytz, H. W., Van Der Geest, R. J., De Koning, P. J., Larsson, H. B., Olesen, J., & Ashina, M. (2012). Headache and prolonged dilatation of the middle meningeal artery by PACAP38 in healthy volunteers. Cephalalgia, 32(2), 140–149. 10.1177/0333102411431333

Amin, F. M., Hougaard, A., Schytz, H. W., Asghar, M. S., Lundholm, E., Parvaiz, A. I., De Koning, P. J. H., Andersen, M. R., Larsson, H. B. W., Fahrenkrug, J., Olesen, J., & Ashina, M. (2014). Investigation of the pathophysiological mechanisms of migraine attacks induced by pituitary adenylate cyclase-activating polypeptide-38. Brain, 137(3), 779–794. 10.1093/brain/awt369

Amin, F. M., & Schytz, H. W. (2018). Transport of the pituitary adenylate cyclase-activating polypeptide across the blood-brain barrier: implications for migraine. The Journal of Headache and Pain, 19(1). 10.1186/s10194-018-0861-3

Arquizan, R., Melchior, A. G., Christensen, R. H., Al-Khazali, H. M., Ashina, M., & Ashina, H. (2025). Vascular responses to molecular migraine triggers: a systematic review of magnetic resonance angiography studies. The Journal of Headache and Pain, 26(1). 10.1186/s10194-025-02105-0

Ashina, H., Christensen, R. H., & Ashina, M. (2022). Provoked versus spontaneous migraine attacks: pathophysiological similarities and differences. The Journal of Headache and Pain, 23(1). 10.1186/s10194-022-01464-2

Ashina, M., Hansen, J. M., Do, T. P., Melo-Carrillo, A., Burstein, R., & Moskowitz, M. A. (2019). Migraine and the trigeminovascular system—40 years and counting. The Lancet Neurology, 18(8), 795–804. 10.1016/s1474-4422(19)30185-1

Ashina, M., Phul, R., Khodaie, M., Löf, E., & Florea, I. (2024). A Monoclonal Antibody to PACAP for Migraine Prevention. New England Journal of Medicine, 391(9), 800–809. 10.1056/nejmoa2314577

Banks, W. A., Kastin, A. J., Komaki, G., & Arimura, A. (1993). Passage of pituitary adenylate cyclase activating polypeptide1-27 and pituitary adenylate cyclase activating polypeptide1-38 across the blood-brain barrier. The Journal of Pharmacology and Experimental Therapeutics, 267(2), 690–696. 10.1016/s0022-3565(25)39457-7

Bates, E., Nikai, T., Brennan, K., Fu, Y. H., Charles, A., Basbaum, A., Ptáček, L., & Ahn, A. (2010). Sumatriptan alleviates nitroglycerin-induced mechanical and thermal allodynia in mice. Cephalalgia, 30(2), 170–178. 10.1111/j.1468-2982.2009.01864.x

Baun, M., Hay-Schmidt, A., Edvinsson, L., Olesen, J., & Jansen-Olesen, I. (2011). Pharmacological characterization and expression of VIP and PACAP receptors in isolated cranial arteries of the rat. European Journal of Pharmacology, 670(1), 186–194. 10.1016/j.ejphar.2011.08.016

Blaeser, A. S., Zhao, J., Sugden, A. U., Carneiro-Nascimento, S., Andermann, M. L., & Levy, D. (2024). Sensitization of meningeal afferents to locomotion-related meningeal deformations in a migraine model. eLife, 12. 10.7554/elife.91871

Christensen, R. H., Strassman, A., Ashina, M., Ashina, H., & Burstein, R. (2025). Activation and sensitization of meningeal nociceptors by PACAP-38: implications for migraine headache. Brain. 10.1093/brain/awaf284

Christensen, S. L., Rasmussen, R. H., Ernstsen, C., La Cour, S., David, A., Chaker, J., Haanes, K. A., Christensen, S. T., Olesen, J., & Kristensen, D. M. (2021). CGRP-dependent signalling pathways involved in mouse models of GTN-cilostazol- and levcromakalim-induced migraine. Cephalalgia, 41(14), 1413– 1426. 10.1177/03331024211038884

Claassen, J. A. H. R., Thijssen, D. H. J., Panerai, R. B., & Faraci, F. M. (2021). Regulation of cerebral blood flow in humans: physiology and clinical implications of autoregulation. Physiological Reviews, 101(4), 1487–1559. 10.1152/physrev.00022.2020

Degel, C., Zitelli, K., Zapata, J., Nassi, J., & Botta, P. (2024). Harnessing Miniscope Imaging in Freely Moving Animals to Unveil Migraine Pathophysiology and Validate Novel Therapeutic Strategies. Synapse, 78(6). 10.1002/syn.70001

Ernstsen, C., Christensen, S. L., Rasmussen, R. H., Nielsen, B. S., Jansen-Olesen, I., Olesen, J., & Kristensen, D. M. (2022). The PACAP pathway is independent of CGRP in mouse models of migraine: possible new drug target? Brain, 145(7), 2450–2460. 10.1093/brain/awac040

Fontaine, D., Almairac, F., Santucci, S., Fernandez, C., Dallel, R., Pallud, J., & Lanteri-Minet, M. (2018). Dural and pial pain-sensitive structures in humans: new inputs from awake craniotomies. Brain, 141(4), 1040–1048. 10.1093/brain/awy005

Gao, Y.-R., & Drew, P. J. (2016). Effects of Voluntary Locomotion and Calcitonin Gene-Related Peptide on the Dynamics of Single Dural Vessels in Awake Mice. The Journal of Neuroscience, 36(8), 2503–2516. 10.1523/jneurosci.3665-15.2016

Gao, Y.-R., Ma, Y., Zhang, Q., Winder, A. T., Liang, Z., Antinori, L., Drew, P. J., & Zhang, N. (2017). Time to wake up: Studying neurovascular coupling and brain-wide circuit function in the un-anesthetized animal. NeuroImage, 153, 382–398. 10.1016/j.neuroimage.2016.11.069

Ghosh, K. K., Burns, L. D., Cocker, E. D., Nimmerjahn, A., Ziv, Y., Gamal, A. E., & Schnitzer, M. J. (2011). Miniaturized integration of a fluorescence microscope. Nature Methods, 8(10), 871–878. 10.1038/nmeth.1694

Gozalov, A., Petersen, K., Mortensen, C., Jansen-Olesen, I., Klaerke, D., & Olesen, J. (2005). Role of K _ATP_ Channels in the Regulation of Rat Dura and Pia Artery Diameter. Cephalalgia, 25(4), 249–260. 10.1111/j.1468-2982.2004.00848.x

Grände, G., Nilsson, E., & Edvinsson, L. (2013). Comparison of responses to vasoactive drugs in human and rat cerebral arteries using myography and pressurized cerebral artery method. Cephalalgia, 33(3), 152–159. 10.1177/0333102412468340

Guo, S., Christensen, S. L., Al-Karagholi, M. A. M., & Olesen, J. (2024). Molecular nociceptive mechanisms in migraine: The migraine cascade. European Journal of Neurology, 31(8). 10.1111/ene.16333

Guo, S., Ernstsen, C., Hay-Schmidt, A., Kristensen, D. M., Ashina, M., Olesen, J., & Christensen, S. L. (2022). PACAP signaling is not involved in GTN- and levcromakalim-induced hypersensitivity in mouse models of migraine. The Journal of Headache and Pain, 23(1). 10.1186/s10194-022-01523-8

Karsan, N., Edvinsson, L., Vecsei, L., & Goadsby, P. J. (2024). Pituitary adenylate cyclase-activating polypeptide targeted treatments in primary headache disorders. Annals of Clinical and Translational Neurology, 11(7), 1654–1668. 10.1002/acn3.52119

Klein, S. P., De Sloovere, V., Meyfroidt, G., & Depreitere, B. (2022). Differential Hemodynamic Response of Pial Arterioles Contributes to a Quadriphasic Cerebral Autoregulation Physiology. Journal of the American Heart Association, 11(1). 10.1161/jaha.121.022943

Kuburas, A., Mason, B. N., Hing, B., Wattiez, A.-S., Reis, A. S., Sowers, L. P., Moldovan Loomis, C., Garcia-Martinez, L. F., & Russo, A. F. (2021). PACAP Induces Light Aversion in Mice by an Inheritable Mechanism Independent of CGRP. The Journal of Neuroscience, 41(21), 4697–4715. 10.1523/jneurosci.2200-20.2021

Lenti, L., Domoki, F., Kis, D., Hegyi, O., Toth, G. K., Busija, D. W., & Bari, F. (2007). Pituitary adenylate cyclase-activating polypeptide induces pial arteriolar vasodilation through cyclooxygenase-dependent and independent mechanisms in newborn pigs. Brain Research, 1165, 81–88. 10.1016/j.brainres.2007.06.032

Levy, D., & Moskowitz, M. A. (2023). Meningeal Mechanisms and the Migraine Connection. Annu Rev Neurosci, 46, 39–58. 10.1146/annurev-neuro-080422-105509

Lund, A. M., & Hannibal, J. (2022). Localization of the neuropeptides pituitary adenylate cyclase-activating polypeptide, vasoactive intestinal peptide, and their receptors in the basal brain blood vessels and trigeminal ganglion of the mouse CNS; an immunohistochemical study. Frontiers in Neuroanatomy, 16. 10.3389/fnana.2022.991403

Masamoto, K., & Kanno, I. (2012). Anesthesia and the Quantitative Evaluation of Neurovascular Coupling. Journal of Cerebral Blood Flow & Metabolism, 32(7), 1233–1247. 10.1038/jcbfm.2012.50

Nilsson, O. R., Kari, L., Rosenke, R., & Steele-Mortimer, O. (2022). Protocol for RNA fluorescence in situ hybridization in mouse meningeal whole mounts. STAR Protocols, 3(2), 101256. 10.1016/j.xpro.2022.101256

Olesen, J. (2024). Provocation of attacks to discover migraine signaling mechanisms and new drug targets: early history and future perspectives - a narrative review. The Journal of Headache and Pain, 25(1). 10.1186/s10194-024-01796-1

Onderwater, G. L. J., Dool, J., Ferrari, M. D., & Terwindt, G. M. (2020). Premonitory symptoms in glyceryl trinitrate triggered migraine attacks: a case-control study. Pain, 161(9), 2058–2067. 10.1097/j.pain.0000000000001894

Pellesi, L., Ashina, M., & Martelletti, P. (2024). Targeting the PACAP-38 pathway is an emerging therapeutic strategy for migraine prevention. Expert Opinion on Emerging Drugs, 29(1), 57–64. 10.1080/14728214.2024.2317778

Perens, J., Salinas, C. G., Skytte, J. L., Roostalu, U., Dahl, A. B., Dyrby, T. B., Wichern, F., Barkholt, P., Vrang, N., Jelsing, J., & Hecksher-Sørensen, J. (2021). An Optimized Mouse Brain Atlas for Automated Mapping and Quantification of Neuronal Activity Using iDISCO+ and Light Sheet Fluorescence Microscopy. Neuroinformatics, 19(3), 433–446. 10.1007/s12021-020-09490-8

Pradhan, A. A., Smith, M. L., McGuire, B., Tarash, I., Evans, C. J., & Charles, A. (2014). Characterization of a novel model of chronic migraine. Pain, 155(2), 269–274. 10.1016/j.pain.2013.10.004

Raffaelli, B., Do, T. P., Chaudhry, B. A., Amin, F. M., Ashina, H., Snellman, J., Maio-Twofoot, T., & Ashina, M. (2024). Activation of ATP-sensitive potassium channels triggers migraine attacks independent of calcitonin gene-related peptide receptors: a randomized placebo-controlled trial. Cephalalgia, 44(1). 10.1177/03331024231222916

Raggi, A., Leonardi, M., Arruda, M., Caponnetto, V., Castaldo, M., Coppola, G., Della Pietra, A., Fan, X., Garcia-Azorin, D., Gazerani, P., Grangeon, L., Grazzi, L., Hsiao, F.-J., Ihara, K., Labastida-Ramirez, A., Lange, K. S., Lisicki, M., Marcassoli, A., Montisano, D. A.,…Martelletti, P. (2024). Hallmarks of primary headache: part 1 – migraine. The Journal of Headache and Pain, 25(1). 10.1186/s10194-024-01889-x

Rakymzhan, A., Li, Y., Tang, P., & Wang, R. K. (2021). Differences in cerebral blood vasculature and flow in awake and anesthetized mouse cortex revealed by quantitative optical coherence tomography angiography. Journal of Neuroscience Methods, 353, 109094. 10.1016/j.jneumeth.2021.109094

Rasmussen, N. B., Deligianni, C., Christensen, C. E., Karlsson, W. K., Al-Khazali, H. M., Van De Casteele, T., Granhall, C., Amin, F. M., & Ashina, M. (2023). The effect of Lu AG09222 on PACAP38- and VIP-induced vasodilation, heart rate increase, and headache in healthy subjects: an interventional, randomized, double-blind, parallel-group, placebo-controlled study. The Journal of Headache and Pain, 24(1). 10.1186/s10194-023-01599-w

Rhea, E. M., Bullock, K. M., & Banks, W. A. (2018). Effect of controlled cortical impact on the passage of pituitary adenylate cyclase activating polypeptide (PACAP) across the blood-brain barrier. Peptides, 99, 8–13. 10.1016/j.peptides.2017.10.013

Schoonman, G. G., Van Der Grond, J., Kortmann, C., Van Der Geest, R. J., Terwindt, G. M., & Ferrari, M. D. (2008). Migraine headache is not associated with cerebral or meningeal vasodilatation—a 3T magnetic resonance angiography study. Brain, 131(8), 2192–2200. 10.1093/brain/awn094

Sokolov, A. Y., Volynsky, M. A., Zaytsev, V. V., Osipchuk, A. V., & Kamshilin, A. A. (2021). Advantages of imaging photoplethysmography for migraine modeling: new optical markers of trigemino-vascular activation in rats. The Journal of Headache and Pain, 22(1). 10.1186/s10194-021-01226-6

Steinmetz, J. D., Seeher, K. M., Schiess, N., Nichols, E., Cao, B., Servili, C., Cavallera, V., Cousin, E., Hagins, H., Moberg, M. E., Mehlman, M. L., Abate, Y. H., Abbas, J., Abbasi, M. A., Abbasian, M., Abbastabar, H., Abdelmasseh, M., Abdollahi, M., Abdollahi, M.,…Dua, T. (2024). Global, regional, and national burden of disorders affecting the nervous system, 1990–2021: a systematic analysis for the Global Burden of Disease Study 2021. The Lancet Neurology, *23*(4), 344–381. 10.1016/s1474-4422(24)00038-3

Syed, A. U., Koide, M., Braas, K. M., May, V., & Wellman, G. C. (2012). Pituitary Adenylate Cyclase-Activating Polypeptide (PACAP) Potently Dilates Middle Meningeal Arteries: Implications for Migraine. Journal of Molecular Neuroscience, 48(3), 574–583. 10.1007/s12031-012-9851-0

Terrier, L. M., Hadjikhani, N., Velut, S., Magnain, C., Amelot, A., Bernard, F., Zöllei, L., & Destrieux, C. (2021). The trigeminal system: The meningovascular complex— A review. Journal of Anatomy, 239(1), 1–11. 10.1111/joa.13413

Tran, C. H. T., & Gordon, G. R. (2015). Acute two-photon imaging of the neurovascular unit in the cortex of active mice. Frontiers in Cellular Neuroscience, 9. 10.3389/fncel.2015.00011

Uddman, R., Goadsby, P. J., Jansen, I., & Edvinsson, L. (1993). PACAP, a VIP-like Peptide: Immunohistochemical Localization and Effect upon Cat Pial Arteries and Cerebral Blood Flow. Journal of Cerebral Blood Flow & Metabolism, 13(2), 291–297. 10.1038/jcbfm.1993.36

Wen, R., Qin, W., Li, B., Li, T., & Xi, L. (2026). Simultaneous mesoscopic imaging of neural activity and hemodynamics reveals anesthetic modulation of neurovascular responses. Photoacoustics, 100874. 10.1016/j.pacs.2026.100874

Wiggers, A., Ashina, H., Hadjikhani, N., Sagare, A., Zlokovic, B. V., Lauritzen, M., & Ashina, M. (2022). Brain barriers and their potential role in migraine pathophysiology. The Journal of Headache and Pain, 23(1). 10.1186/s10194-021-01365-w

Witztum, J., Singh, A., Zhang, R., Johnson, M., & Liston, C. (2023). An automated platform for Assessing Working Memory and prefrontal circuit function. Neurobiology of Stress, 24, 100518. 10.1016/j.ynstr.2023.100518

