## Supplemental Figure 1 for "High spatiotemporal resolution in vivo imaging reveals preferential PACAP38 action on dural vasculature and a functional therapeutic readout"

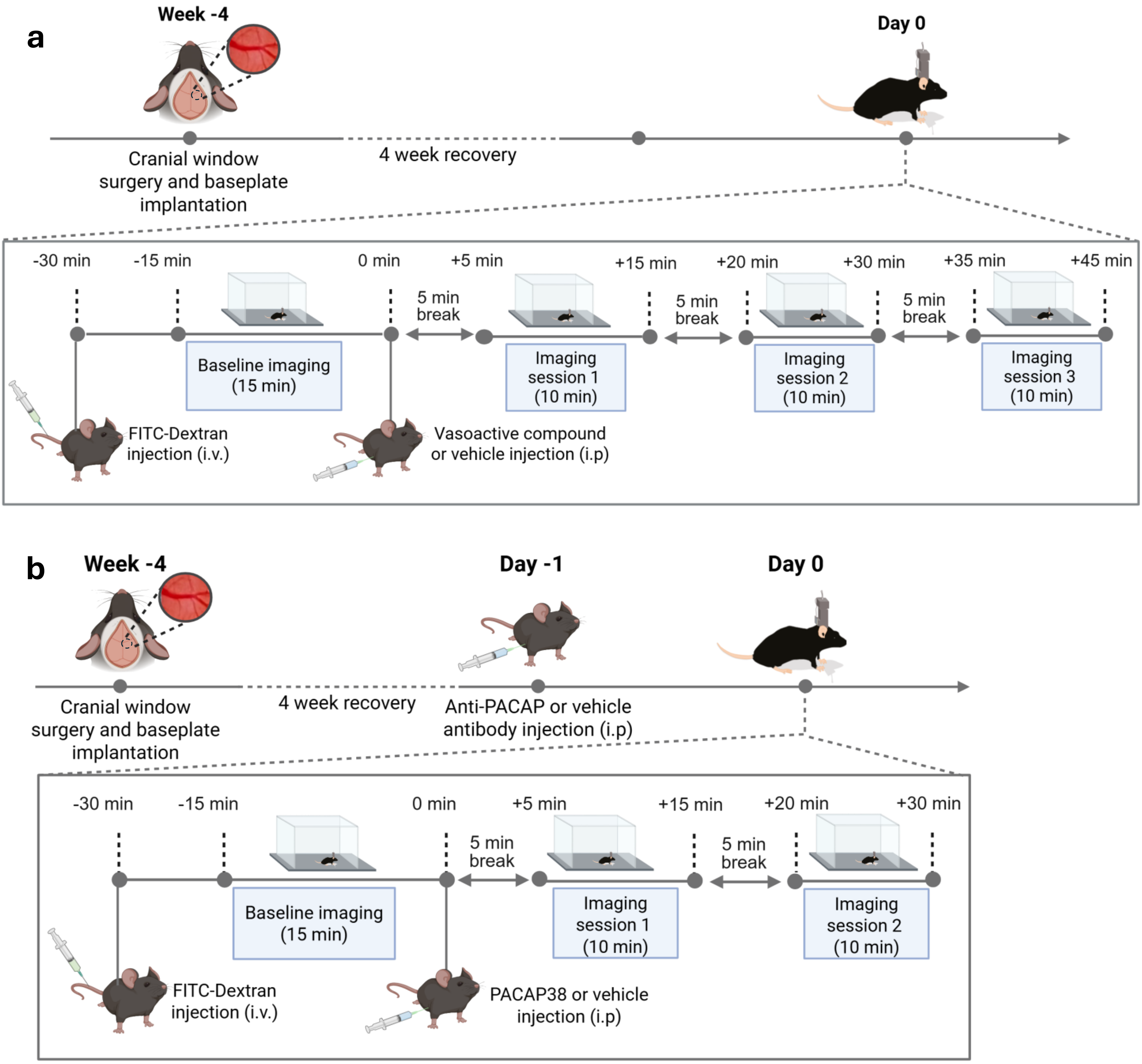

a
Week -4
Day 0
Cranial window surgery and baseplate implantation
4 week recovery
-30 min
-15 min
0 min
+5 min
+15 min
+20 min
+30 min
+35 min
+45 min
5 min break
5 min break
5 min break
Baseline imaging (15 min)
Imaging session 1 (10 min)
Imaging session 2 (10 min)
Imaging session 3 (10 min)
FITC-Dextran injection (i.v.)
Vasoactive compound or vehicle injection (i.p)
b
Week -4
Day -1
Day 0
Cranial window surgery and baseplate implantation
4 week recovery
Anti-PACAP or vehicle antibody injection (i.p)
-30 min
-15 min
0 min
+5 min
+15 min
+20 min
+30 min
5 min break
5 min break
Baseline imaging (15 min)
Imaging session 1 (10 min)
Imaging session 2 (10 min)
FITC-Dextran injection (i.v.)
PACAP38 or vehicle injection (i.p)
