## Supplemental Figure 2 for "High spatiotemporal resolution in vivo imaging reveals preferential PACAP38 action on dural vasculature and a functional therapeutic readout"

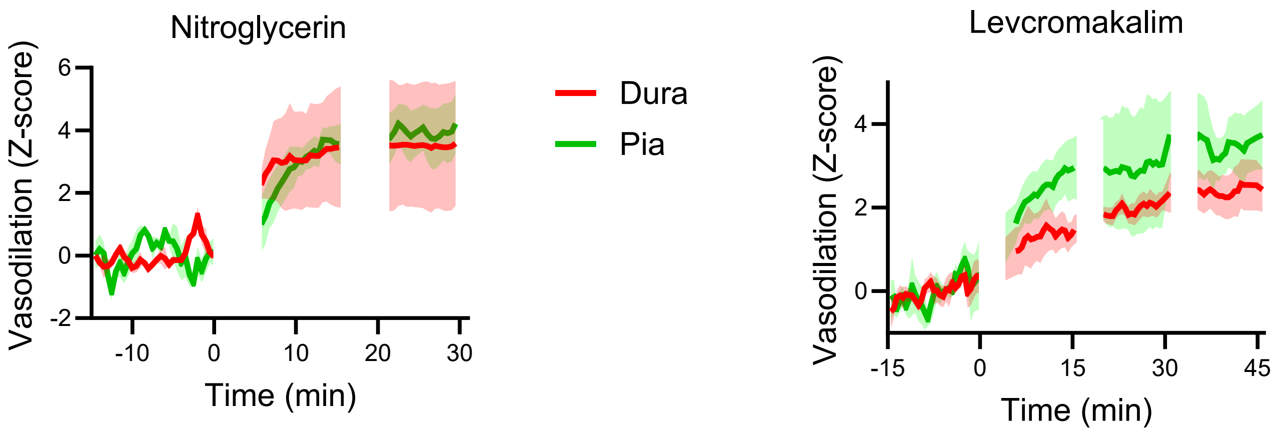
Nitroglycerin
Vasodilation (Z-score)
6
4
2
0
-2
-10
0
10
20
30
Time (min)
Dura
Pia
Levcromakalim
Vasodilation (Z-score)
4
2
0
-15
0
15
30
45
Time (min)
