## Supplemental Figure 3 for "High spatiotemporal resolution in vivo imaging reveals preferential PACAP38 action on dural vasculature and a functional therapeutic readout"

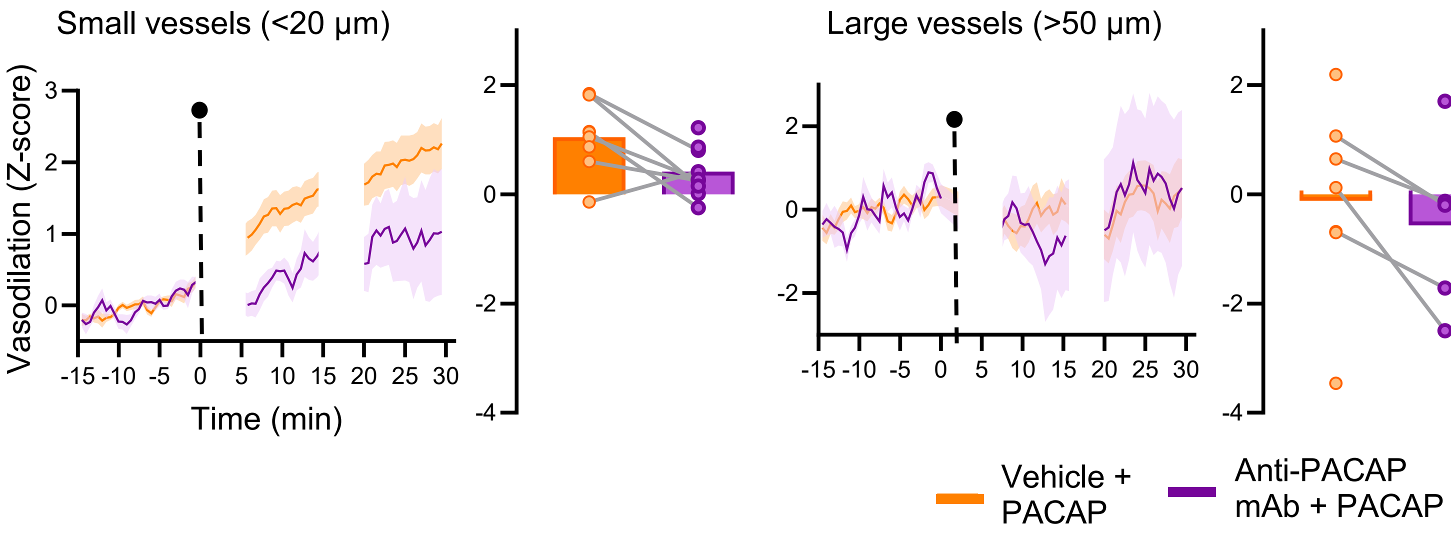

Small vessels (<20 μm)
Large vessels (>50 μm)
Vasodilation (Z-score)
Time (min)
-15 -10 -5 0 5 10 15 20 25 30
Vehicle + PACAP
Anti-PACAP mAb + PACAP
