## Supplemental Figure 4 for "High spatiotemporal resolution in vivo imaging reveals preferential PACAP38 action on dural vasculature and a functional therapeutic readout"

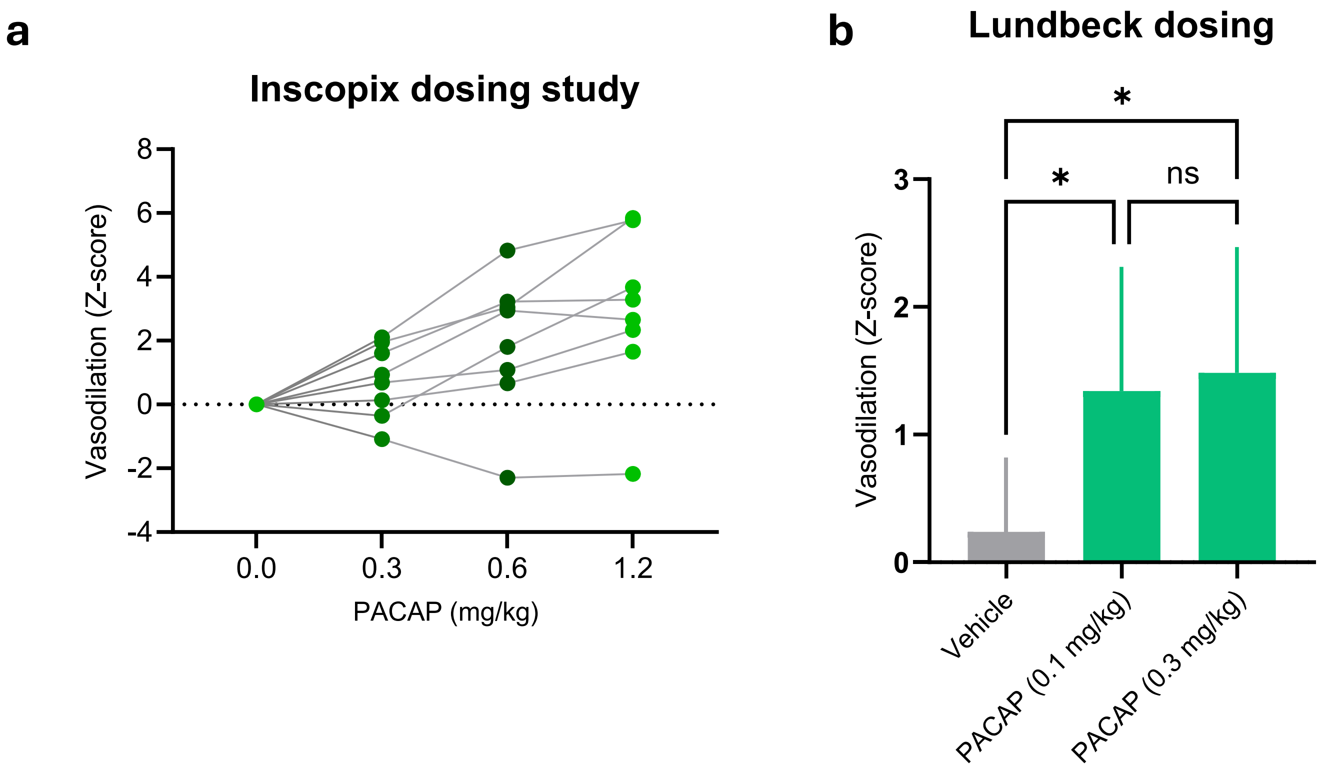

a
Inscopix dosing study
Vasodilation (Z-score)
8
6
4
2
0
-2
-4
0.0
0.3
0.6
1.2
PACAP (mg/kg)
b
Lundbeck dosing
*
*
ns
Vasodilation (Z-score)
3
2
1
0
Vehicle
PACAP (0.1 mg/kg)
PACAP (0.3 mg/kg)
