## Supplemental Table 1 for "High spatiotemporal resolution in vivo imaging reveals preferential PACAP38 action on dural vasculature and a functional therapeutic readout"

| **Treatment** | **Vessel category** | **n** | **Z-score** |  | **Percent change (%)** |
| --- | --- | --- | --- | --- | --- |
| **Levcromakalim study — Figs. 1–2 and Fig. S2** | | | | | |
| Levcromakalim, 1 mg/kg | All vessels | 7 | 1.61 ± 0.38 |  | 7.01 ± 1.45 |
| Levcromakalim, 1 mg/kg | Small (<20 µm) | 7 | 1.48 ± 0.42 |  | 7.34 ± 1.93 |
| Levcromakalim, 1 mg/kg | Large (>50 µm) | 5 | 3.03 ± 1.00 |  | 6.19 ± 2.28 |
| Levcromakalim, 1 mg/kg | Dura | 4 | 1.32 ± 0.53 |  | 6.17 ± 2.16 |
| Levcromakalim, 1 mg/kg | Pia | 3 | 2.45 ± 0.59 |  | 7.33 ± 1.90 |
| **Nitroglycerin study — Figs. 1–2 and Fig. S2** | | | | | |
| Nitroglycerin, 10 mg/kg | All vessels | 9 | 1.93 ± 0.26 |  | 5.40 ± 0.58 |
| Nitroglycerin, 10 mg/kg | Small (<20 µm) | 9 | 1.66 ± 0.34 |  | 5.30 ± 0.77 |
| Nitroglycerin, 10 mg/kg | Large (>50 µm) | 7 | 1.90 ± 0.72 |  | 2.05 ± 0.95 |
| Nitroglycerin, 10 mg/kg | Dura | 2 | 3.07 ± 1.47 |  | 10.03 ± 2.16 |
| Nitroglycerin, 10 mg/kg | Pia | 3 | 2.77 ± 0.59 |  | 6.53 ± 2.37 |
| **PACAP38 acute study — Figs. 1–2 and Fig. S4b** | | | | | |
| PACAP38, 0.1 mg/kg | All vessels | 11 | 1.34 ± 0.29 |  | 5.78 ± 1.25 |
| PACAP38, 0.1 mg/kg | Small (<20 µm) | 11 | 1.47 ± 0.36 |  | 6.76 ± 1.62 |
| PACAP38, 0.1 mg/kg | Large (>50 µm) | 6 | -0.32 ± 0.49 |  | -0.67 ± 0.83 |
| **PACAP38 compartment analysis — Fig. 3h** | | | | | |
| PACAP38, 0.1 mg/kg | Dura | 10 | 3.961 ± 0.915 |  | 18.74 ± 4.72 |
| PACAP38, 0.1 mg/kg | Pia | 7 | -0.48 ± 0.62 |  | -0.92 ± 1.27 |
| **Anti-PACAP pretreatment study — Fig. 4 and Fig. S3** | | | | | |
| Vehicle + PACAP38 | All vessels | 9 | 0.88 ± 0.25 |  | 2.65 ± 0.81 |
| Vehicle + PACAP38 | Small (<20 µm) | 9 | 1.19 ± 0.25 |  | 3.80 ± 0.93 |
| Vehicle + PACAP38 | Large (>50 µm) | 7 | -0.12 ± 0.68 |  | 0.49 ± 1.03 |
| Vehicle + PACAP38 | Dura | 5 | 5.31 ± 1.82 |  | 17.96 ± 4.21 |
| Vehicle + PACAP38 | Pia | 6 | -0.70 ± 0.69 |  | -1.31 ± 1.44 |
| Anti-PACAP mAb + PACAP38 | All vessels | 9 | 0.30 ± 0.13 |  | 0.63 ± 0.35 |
| Anti-PACAP mAb + PACAP38 | Small (<20 µm) | 9 | 0.41 ± 0.15 |  | 0.96 ± 0.44 |
| Anti-PACAP mAb + PACAP38 | Large (>50 µm) | 5 | -0.57 ± 0.73 |  | -1.90 ± 2.66 |
| Anti-PACAP mAb + PACAP38 | Dura | 5 | 1.32 ± 0.63 |  | 3.18 ± 1.17 |
| Anti-PACAP mAb + PACAP38 | Pia | 4 | -0.53 ± 0.31 |  | -1.08 ± 0.48 |
